# Reverse engineering the fatally cross-reactive A3A TCR to decouple potency and specificity

**DOI:** 10.64898/2026.05.03.722528

**Authors:** Julia V. McCarthy, A. Christina Heroven, King Ifashe, Martin Fellermeyer, Dan Hudson, Raul Cioaca, Max N. Quastel, Yunkai Yang, Matteo Cagiada, Simeon D. Draganov, Christopher J. Thorpe, Adán Pinto-Fernández, Alexander Greenshields-Watson, Geraldine M. Gillespie, Charlotte M. Deane, Ricardo A. Fernandes

## Abstract

T cell receptor (TCR) affinity enhancement can introduce off-target cross-reactivity with life-threatening consequences, as illustrated by the MAGE-A3-specific A3A TCR, which caused fatal cardiotoxicity through recognition of a Titin-derived peptide. Here, we reconstructed the cross-reactivity landscape by reverse-engineering A3A toward its wild-type precursor, generating intermediate variants in which engineered CDR2α residues are systematically reverted to the wild-type sequence. Reverting just two engineered residues yields a receptor, v9, that retains MAGE-A3 cytotoxicity comparable to A3A while eliminating Titin and other acquired cross-reactivities. Structurally, these substitutions reduce CDR2α-MHC contacts and disrupt an intra-TCR CDR2α-CDR3β interaction, propagating conformational changes across CDR3 loops that reshape peptide engagement without altering docking geometry. These results demonstrate that mutations outside the peptide-contacting CDR3 loops can allosterically reconfigure antigen specificity and establish simple stepwise reverse engineering to wild-type as a strategy for correcting TCR cross-reactivity.

## Introduction

T cell receptor (TCR) affinity enhancement is a central strategy for boosting immune responses against tumour antigens^1–3^, including self-derived, low-affinity targets such as melanoma-associated antigens (MAGE) that evade thymic selection and are thus relatively common targets in many cancers^4^. However, engineered potency carries an inherent risk: affinity-enhanced TCRs can acquire unpredictable cross-reactivities with off-target self-antigens, leading to severe toxicities^5,6^. A prominent example is the A3A TCR, an affinity-enhanced derivative of the wild-type MAGE-A3-specific A3 TCR isolated from a melanoma patient^7,8^. The A3A differs from its precursor by only four CDR2α residues, yet this was sufficient to cause fatal cardiotoxicity through recognition of a Titin-derived peptide expressed on cardiac myocytes^7^. Enhancing TCR affinity has been achieved repeatedly, however engineering both affinity and specificity remains a challenge and a major limitation for the development of safe TCR-T therapeutics. These challenges highlight a fundamental gap in our understanding of how minimal sequence changes in engineered TCRs reshape peptide recognition and introduce cross-reactivity.

Efforts to map and predict TCR specificity have advanced considerably in recent years. High-throughput technologies such as peptide-MHC (pMHC) yeast-display^9–11^, genome-wide functional screens (T-Scan)^12^, and synthetic circuit-based platforms (TCR-MAP)^13^ have revealed the extensive degeneracy of TCR recognition, demonstrating that small sequence changes can produce broad and unexpected cross-reactivity. However, most analyses have examined TCRs at single endpoints; either natural precursors or engineered high-affinity derivatives, without capturing the trajectory linking them^14^. Moreover, mechanistic understanding remains centred around CDR3 loops, long viewed as the dominant determinants of peptide recognition^15–18^. Far less is known about the contribution of non-CDR3 loops to peptide recognition, and whether mutations in these regions, particularly those contacting the MHC rather than the peptide, can be used to reshape specificity.

This question can be addressed by reversing the conventional engineering trajectory. Thymically selected TCRs have passed central tolerance checkpoints that constrain self-reactivity^19–21^, and their wild-type sequences therefore capture an inherent safety profile that can be lost during affinity maturation. We hypothesised that systematically reverting engineered residues to the parental sequence could reduce cross-reactivity and offer a simple alternative for rescuing clinically developed but toxic TCRs. The A3A/A3 WT pair, differing by only four CDR2α residues, provides an ideal system to test this because the mutational space is constrained enough to be systematically explored.

Here, we generate a panel of eight intermediate TCR variants spanning 8 of the 14 possible intermediate CDR2α sequences between A3 WT and A3A, and characterise each TCR using large-scale pMHC yeast-display screening, *in vitro* co-culture and primary T cell cytotoxic assays, X-ray crystallography, and all-atom molecular dynamics (MD) simulations. We identify v9, a variant in which two of the four engineered residues are reverted to wild-type, as a receptor that matches the cytotoxicity of A3A against MAGE-A3-expressing targets while eliminating Titin cross-reactivity. Structural and MD analyses reveal that the CDR2α substitutions, which largely contact the MHC and not the peptide, propagate allosterically to reconfigure CDR3 conformational ensembles and reshape peptide contacts, providing a mechanistic explanation for how non-CDR3 mutations control peptide specificity. Notably, cross-reactivities intrinsic to the wild-type precursor persisted across all variants, indicating that the parental receptor sets a reactivity baseline that this approach cannot easily overcome. These findings establish stepwise reversion to wild-type as a general strategy for both dissecting and correcting cross-reactivity. Such exercises may have direct translational applications but also serve to inform our understanding of fundamental TCR recognition rules.

## Results

### Tracing the stepwise evolution of cross-reactivity

The engineered A3A TCR differs from its precursor, the wild-type A3 TCR (A3 WT)^8^, by only four CDR2α residues, and yet these were sufficient to increase its affinity to MAGE-A3 by approximately 250-fold and introduce self-reactivity to Titin^7^. To define how the individual CDR2α residues contribute to sensitivity and specificity, we generated a panel of intermediate TCR variants in which one, two, or three of the mutated A3A CDR2α residues (LVRPY) were reverted to the original A3 WT sequence (LIQSS) (**Fig. 1a,b**). We reasoned that by restricting the re-engineering of the CDR2α to the boundaries set by the A3 WT and A3A, it would be possible to identify variants that re-acquire the original safety profile of the A3 TCR without compromising on MAGE-A3 reactivity (**Fig. 1a)**. We primarily focused our efforts on residues 52-54 given their greater proximity to the MHC compared to residue 51, as observed in the structure of the A3A-like MAG-IC3 TCR in complex with MAGE-A3-HLA-A*01:01^22^ (**Extended Data Fig. 1a**). To compare peptide recognition profiles across all 10 TCRs (A3 WT, A3A, and eight intermediate variants), we produced and screened each TCR against a yeast-display 9mer-HLA-A*01:01 library comprising millions of unbiased 9-mer peptides (**Extended Data Fig. 1b,c; Extended Data Fig. 2**)^9–11,23^. The library was designed based on the preferences of endogenously expressed HLA-A*01:01 peptides, whereby anchor residues were fixed (Asp/Glu at P3 and Tyr at P9) to maximise peptide stability in the MHC groove (**Extended Data Fig. 1b**). All other positions were randomly assigned a balanced distribution of all amino acid codons (excluding Cys), achieving a peptide diversity of 1.8 × 10^9^. To first confirm correct folding and display of the 9mer-HLA-A*01:01 complexes on the yeast cells^24^, we tested whether A3A could bind to a single MAGE-A3 clone presented by HLA-A*01:01 in a single-chain trimer format (**Extended Data Fig. 1d,e**).

**Fig. 1.**
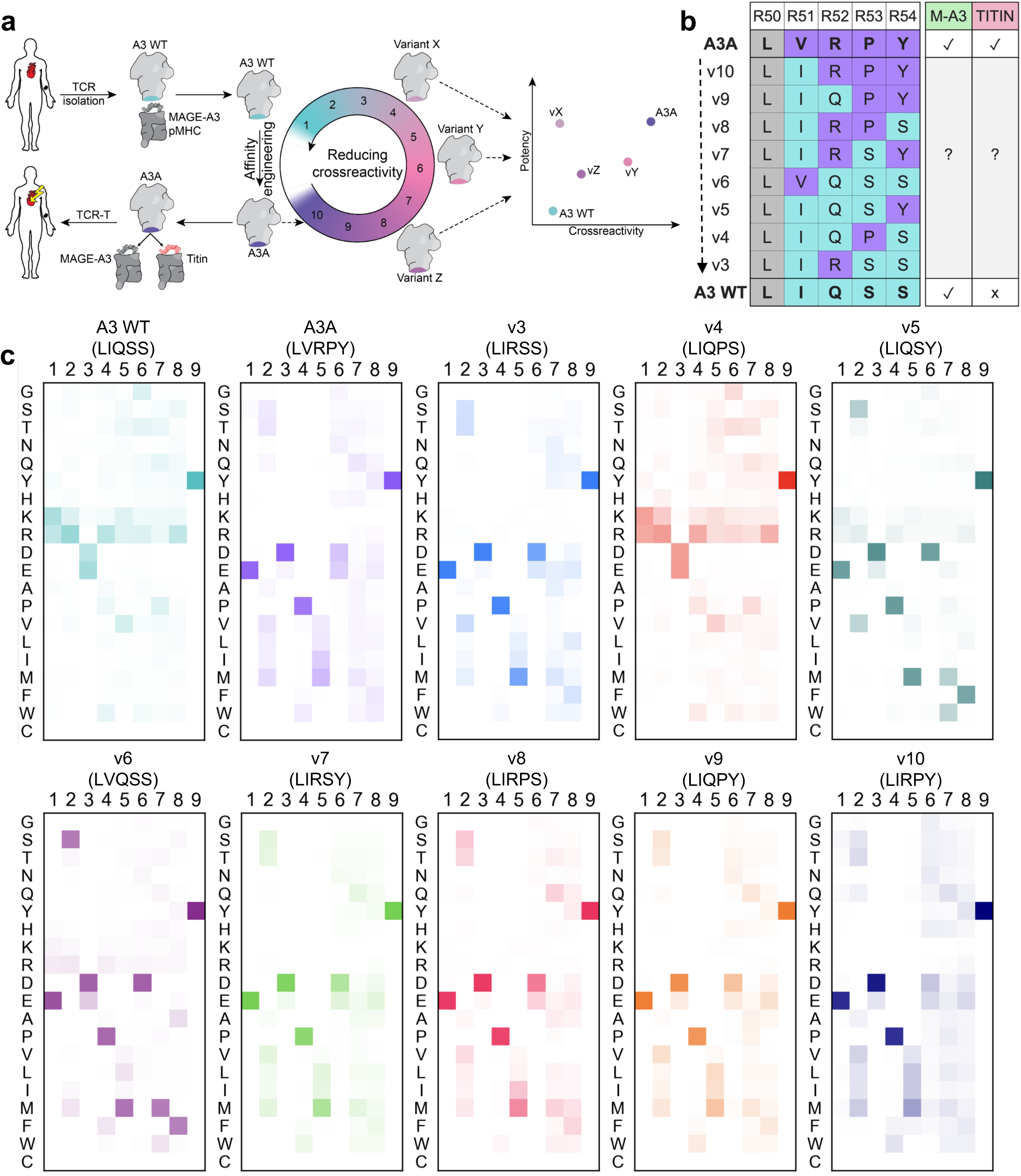
Reverse-engineering the A3A TCR to trace the stepwise evolution of cross-reactivity. **a**, Schematic of the strategy taken to investigate the role of the mutated A3A CDR2α residues on potency (sensitivity) and cross-reactivity (specificity). **b,** Overview of the process of reverting the CDR2α sequence of the A3A to the A3 WT. In descending order, the displayed CDR2α sequences are of the A3A TCR (purple), the eight intermediate TCRs, and the A3 WT TCR (teal). R = residue. Shared amino acids with A3A are in purple and with A3 WT in teal. The R50 residue (L = Leu) shown in grey is shared by both A3A and A3 WT (and thus also by all intermediates). The key on the right indicates the known reactivities of A3 WT and A3A for M-A3 (MAGE-A3; green) and Titin (red), whereby a tick indicates reactivity, a cross indicates no reactivity, and a “?” indicates unknown reactivity. **c,** Positional frequency heatmaps showing the amino acid composition of the peptides enriched after the fourth round of selection with TCRs A3 WT (teal), A3A (purple), and all intermediates (multi-coloured). A darker colour represents a greater frequency of a given amino acid at a specific peptide position. Single-letter abbreviations for the amino acid residues are as follows: A, Ala; C, Cys; D, Asp; E, Glu; F, Phe; G, Gly; H, His; I, Ile; K, Lys; L, Leu; M, Met; N, Asn; P, Pro; Q, Gln; R, Arg; S, Ser; T, Thr; V, Val; W, Trp; and Y, Tyr.

After three and four rounds of selection, the majority of the TCR variants showed robust tetramer staining of enriched yeast-pMHC populations (**Extended Data Fig. 3a,b**), and deep sequencing of the enriched peptides revealed a conserved N-terminal EXDPI motif (**Fig. 1c**) consistent with previous analyses of the A3A’s binding preferences^7,12,22,25^. In contrast, the parental A3 WT failed to enrich a clear peptide motif or achieve robust tetramer staining, suggesting limited peptide cross-reactivity and consistent with its low affinity for MAGE-A3 (*K*_D_ ≈ 500 μM*^7^*). Limited peptide enrichment was additionally observed by v4, while v5 and v6 exhibited only modest enrichment, also indicative of low-affinity interactions. These three low-binding variants share a common feature: retention of A3 WT’s Gln52 together with at least one of the WT-derived Ser residues at 53 or 54 (**Fig. 1b**).

To capture the similarities and differences across the large number of peptide sequences enriched per TCR, we used the peptide:MHC binding energy covariance (PMBEC) amino acid similarity matrix to cluster the top 95% of peptides based on chemical similarity (**Extended Data Fig. 4**) and amino acid preference (**Fig. 2a**)^26^. These analyses revealed robust N-terminal constraints for A3A and TCR variants v3, v7, v8, v9 and v10. To identify peptide recognition hotspots and quantify positional amino acid promiscuity, we next calculated enrichment-normalised average PMBEC-similarity scores at each peptide position, followed by a global average similarity score across the entire peptide for each TCR (**Fig. 2b; Extended Data Fig. 4**). This enabled ranking of all TCR variants by the breadth of their peptide recognition landscapes, providing a quantitative measure of peptide degeneracy, and in turn, specificity. Using A3A as a benchmark (score 6.48), the TCR variants v3, v7, v8 and v9 maintained improved specificity profiles (scores 7.51, 6.99, 7.74 and 7.03, respectively). This was not the case for v10 (score 5.62), explained by the increased degree of amino acid promiscuity at its C-terminus. Notably, enhanced specificity correlated with the presence of Arg52 and at least one Ser53 or Ser54 in the CDR2α loop, except in the case of v9, which combined the A3 WT’s Gln52 with the Pro53 and Tyr54 from the affinity-enhanced A3A sequence.

**Fig. 2.**
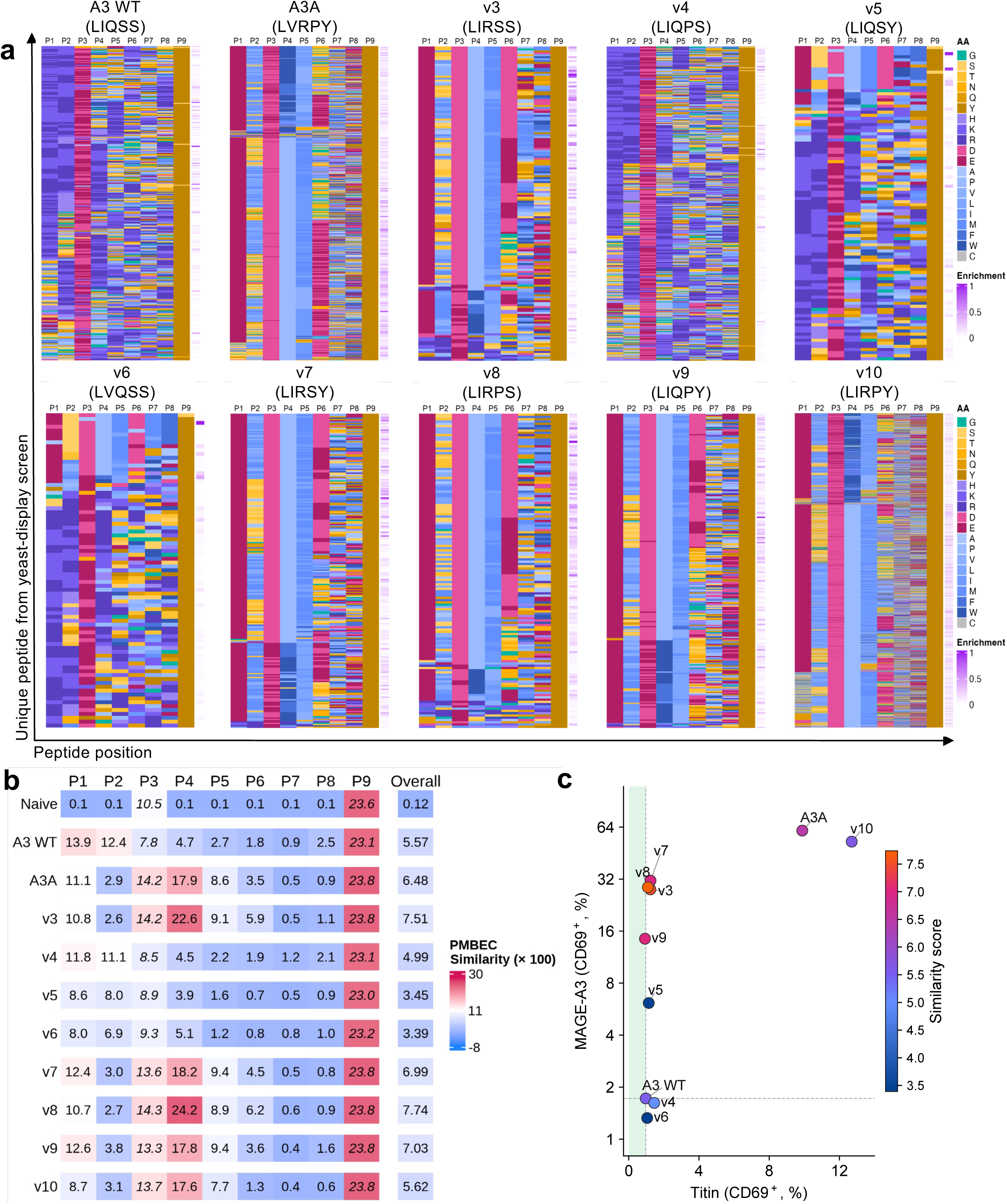
Reactivity to Titin can be decoupled from MAGE-A3. **a**, Heatmaps comparing the top 95% of enriched peptides at round 4 from the pMHC yeast-display sequences. For each individual peptide, the PMBEC score was calculated per position (P1-P9), and the peptides were grouped by hierarchical clustering. Each amino acid was assigned a colour (grouped by chemical similarity; “AA” key), with darker colours representing larger amino acids. The peptide enrichment level is displayed in purple. **b,** PMBEC similarity scores for each peptide position were calculated by taking an enrichment-normalised average of the top 95% of enriched peptides at round 4, excluding the anchor residues (italicised), and multiplying by 100. “Naive” refers to the 9-mer-A*01:01 library prior to TCR screening. Scores correspond to the heatmaps in (Fig. S4). **c,** The CD69 expression levels (%) of the ten A3A-variant-Jurkat T cell lines to Titin vs MAGE-A3 (100 μM) are plotted (corresponding to the data shown in Fig. S5c) and colour-coded to the PMBEC similarity scores (as shown in Fig. 2b).

### Reverse engineering A3A to decouple Titin from MAGE-A3 reactivity

Next, to validate the similarity scores and investigate whether cross-reactivity to Titin was intrinsically linked to robust reactivity to MAGE-A3, all TCR variants were expressed on the surface of CD8⁺ Jurkat T cells and tested for their reactivity to MAGE-A3 and Titin in co-culture assays (**Extended Data Fig. 5a-b**). While most variants retained enhanced sensitivity to MAGE-A3 relative to A3 WT, sensitivity to Titin was strongly diminished in all but a single variant, v10 (LIRPY) (**Fig. 2c** and **Extended Data Fig. 5c**). This was in keeping with v10’s reduced overall specificity score relative to A3A (**Fig. 2b**). In addition, we found the Titin vs MAGE-A3 activation data to be negatively correlated with the specificity scores (**Extended Data Fig. 5d;** r = -0.8667, p = 0.0022), whereby the highest scorers (v3, v7, v8 and v9; **Fig. 2b**) clustered together as the most ideal balance of sensitivity and specificity to MAGE-A3. Taken together, peptide degeneracy analyses and co-culture activation data indicated that variants v3, v7, v8 and v9 represented preferable solutions to A3A, and that subtle mutations in the CDR2α loop could functionally decouple Titin from MAGE-A3 reactivity.

Closer analysis of the CDR2α sequences indicated that the RPY motif was required for recognition of both MAGE-A3 and Titin antigens, as substitution to RPS or RSY eliminated Titin reactivity (**Fig. 2c, Extended Data Fig. 5c,d**). In contrast, sensitivity to MAGE-A3 depended primarily on the presence of a positively charged Arg52, with v9 being the only outlier in this respect (LIQPY). This was most clearly illustrated by comparison of the A3 WT (LIQSS) with v3 (LIRSS), where substitution of a single residue, corresponding to a difference of only six atoms, generated a potent MAGE-A3-reactive TCR.

Next, we used the mimotopes selected by the A3A TCR in the yeast-display pMHC library screen to test whether an *in silico* motif-similarity analysis could predict the known agonist peptides of the A3A TCR^27^. The top 55 candidate antigens out of more than 10 million ranked 9-mer peptides from the human peptidome (**Extended Data Fig. 6a; Supplementary Table 4**) were tested in co-culture assays.

This small set of peptides already contained 9 of the previously validated agonist peptides, including MAGE-A3, -A6, -B18, Titin, and a new additional peptide that had not been detected before, Dynactin subunit 4. Using this set of agonist peptides, we then refined the *in silico* motif similarity analysis to generate new predictions that identified two additional weak agonists, ANKRD16 and FXYD6 (**Extended Data Fig. 6b,c**; **Supplementary Table 5**), amounting to 12 wild-type peptides. The considerable sequence heterogeneity between these agonist peptides illustrates the challenge of predicting TCR agonists, and the degeneracy of TCR/pMHC interactions. Indeed, hFat2 and PLD5 share only three amino acids with MAGE-A3, and yet these peptides triggered stronger activation than MAGE-B18 despite this peptide sharing four additional amino acids with MAGE-A3 (**Extended Data Fig. 6b,c**).

Having found that robust reactivity to MAGE-A3 was not intrinsically coupled to Titin cross-reactivity, we next compared the cross-reactivity profiles of all the TCR variants against the set of 12 wild-type peptides recognised by A3A (**Fig. 3a; Extended Data Fig. 6b**). In agreement with the enrichment profiles in the yeast-display library screen, A3 WT, v4, v5 and v6 showed weak or no responses to this set of antigens. v10 (LIRPY) was as sensitive and cross-reactive as A3A (LVRPY), and v6 (LVQSS) was as weak and specific as A3 WT (LIQSS), confirming that Val51 and Ile51 were not contributing to antigen potency or reactivity and justifying our focus on CDR2α residues 52-54. Importantly, v8 and v9 showed robust reactivity to MAGE-family tumour antigens, but no response to Titin and negligible responses to the other A3A off-target antigens. Of these, v9 had the strongest preference for MAGE antigens, producing only weak responses to the non-MAGE family antigens native to the A3 WT, hFat2 and PLD5^12^.

**Fig. 3.**
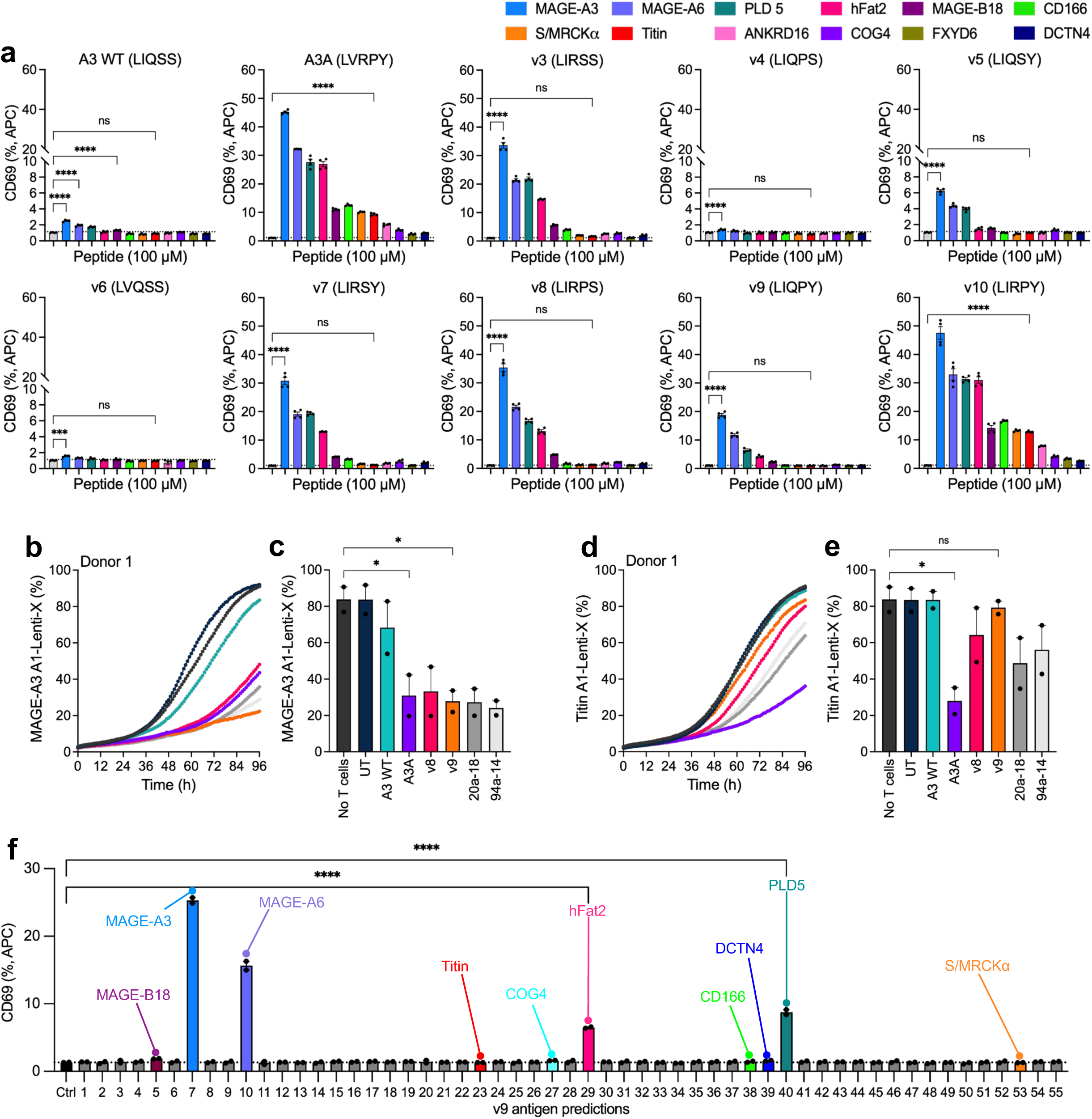
An intermediate TCR is as specific as the A3 WT and as sensitive as the affinity-enhanced A3A. **a**, A1-K562s were pulsed with 100μM of the 12 A3A-agonist peptides (Extended Data Fig. 6b), or a control non-agonist peptide (grey bar, dotted line), and used to stimulate the ten CD8+ A3A-variant-Jurkat T cell lines at an E/T ratio of 1:1 for 18h. Surface CD69 expression was quantified by flow cytometry. Error bars represent means ± SEM of technical replicates pooled from two independent experiments. Statistical analysis by ordinary one-way ANOVA: ****p≤0.0001 ***p≤0.001, ns p>0.05. **b-e,** Primary CD8+ T cells isolated from “donor 1” were transduced with the TCR variants A3 WT, A3A, v8, v9 and two previously engineered A3A-variants 20a-18 and 94a-14 (Zhao et al., Science, 2022). Lenti-X cells were transduced with HLA-A*01:01 (A1-Lenti-X), pulsed with 50 μM of **(b,c)** MAGE-A3 or **(d,e)** Titin peptides, for 1h, and used to stimulate the A3A-variant CD8+ T cells at an E/T ratio of 1:2 for 3 days (UT = untransduced T cells). A1-Lenti-X confluence was monitored by an Incucyte live cell imaging microscope. The cytotoxicity data from “donor 1” were pooled with the cytotoxicity data from a second experiment with “donor 2” (Extended Data Fig. 7a,b), such that **(c,e)** represents quantification of data shown in **(b,d)**. Statistical analysis by ordinary one-way ANOVA: *p≤0.05, ns p>0.05. Error bars represent means of biological duplicates ± SEM. **f,** A1-K562s were pulsed with 100μM of the top 55 predicted agonist peptides of v9 (determined by an *in silico* motif similarity analysis), and a control non-agonist peptide (black bar, dotted line). This library of peptide-pulsed cells was then used to stimulate v9-Jurkat T cells at an E/T ratio of 1:1 for 18h. Anti-CD69 staining was then performed on the v9-Jurkat T cells and CD69 levels were measured by flow cytometry. Error bars represent means of technical duplicates ± SEM, and the data are representative of two independent experiments. Statistical analysis by ordinary one-way ANOVA, ****p≤0.0001. Coloured bars are the known A3A agonists (Extended Data Fig. 6b).

To further validate these reactivity profiles, we performed cytotoxicity assays using primary CD8+ T cells transduced with TCRs A3 WT, A3A, v8 and v9 that were cultured with cells expressing MAGE-A3 and Titin peptides. v9 was the only TCR to exhibit specific cytotoxicity to the MAGE-A3 peptide (**Fig. 3b-e; Extended Data Fig. 7a,b**) and was superior to two additional A3A TCR variants engineered using an alternative catch-bond screening approach^28^. In contrast to Jurkat activation assays (**Fig. 3a**), v8 and v9 expressed in primary CD8+ T cells killed MAGE-A3-pulsed targets at levels comparable to A3A (**Fig. 3b,c; Extended Data Fig. 7a**), including towards endogenously presented MAGE-A3 by the A375 cell line (**Extended Data Fig. 7c-e**). By SPR, we found that v9 showed no detectable binding to Titin (up to 100 𝜇M of the TCR as analyte, at 25°C), while v8 showed a very modest increase in RU, insufficient to determine a *K_D_* but suggestive of potentially very weak binding to this peptide (**Extended Data Fig. 8e-h**). In addition, compared to A3A, binding affinities of v8 and v9 to MAGE-A3 were found to be 9-fold and 15-fold lower, respectively (**Extended Data Fig. 8a-d;** v8/MAGE-A3 *K*_D_ = 13.58 μM, v9/MAGE-A3 *K*_D_ = 23.87 μM).

So far, v9 possessed the most ideal balance of specificity and sensitivity to MAGE antigens. To exclude additional cross-reactivities against a broader set of off-target peptides introduced during its engineering process, we applied the same *in silico* motif similarity analysis that was applied to A3A’s yeast-display library screen to v9’s screen^27^. As with A3A, the top 55 predicted wild-type candidates (**Extended Data Fig. 9a; Supplementary Table 6**) were investigated for activation, and no additional agonist antigens were identified (**Fig. 3f**). In addition, no alloreactivity was observed beyond HLA-A*01:01 to seven of the most common MHC class I alleles (**Extended Data Fig. 9b-e**). Collectively, v9 retained the specificity of the A3 WT while exhibiting the sensitivity of the affinity-enhanced A3A.

### Structural comparison of the A3A and v9 TCR/MAGE-A3-HLA-A*01:01 complexes

To understand the structural impact of reversing the mutated CDR2α residues, we determined the crystal structures of the v9 and A3A TCRs in complex with MAGE-A3-HLA-A*01:01. The complexes diffracted to 2.4 Å (v9 TCR/pMHC) and 2.8 Å (A3A TCR/pMHC) (**Supplementary Table 1**) and displayed largely similar structures and interface properties (**Fig. 4a-c**; r.m.s.d. 0.26 Å). No significant differences were observed between TCR docking angles (56.9° for v9 and 55.9° for A3A), incident angles (13.5° for v9 and 13.6° for A3A), or TCR shifts (**Supplementary Table 2**).

**Fig. 4.**
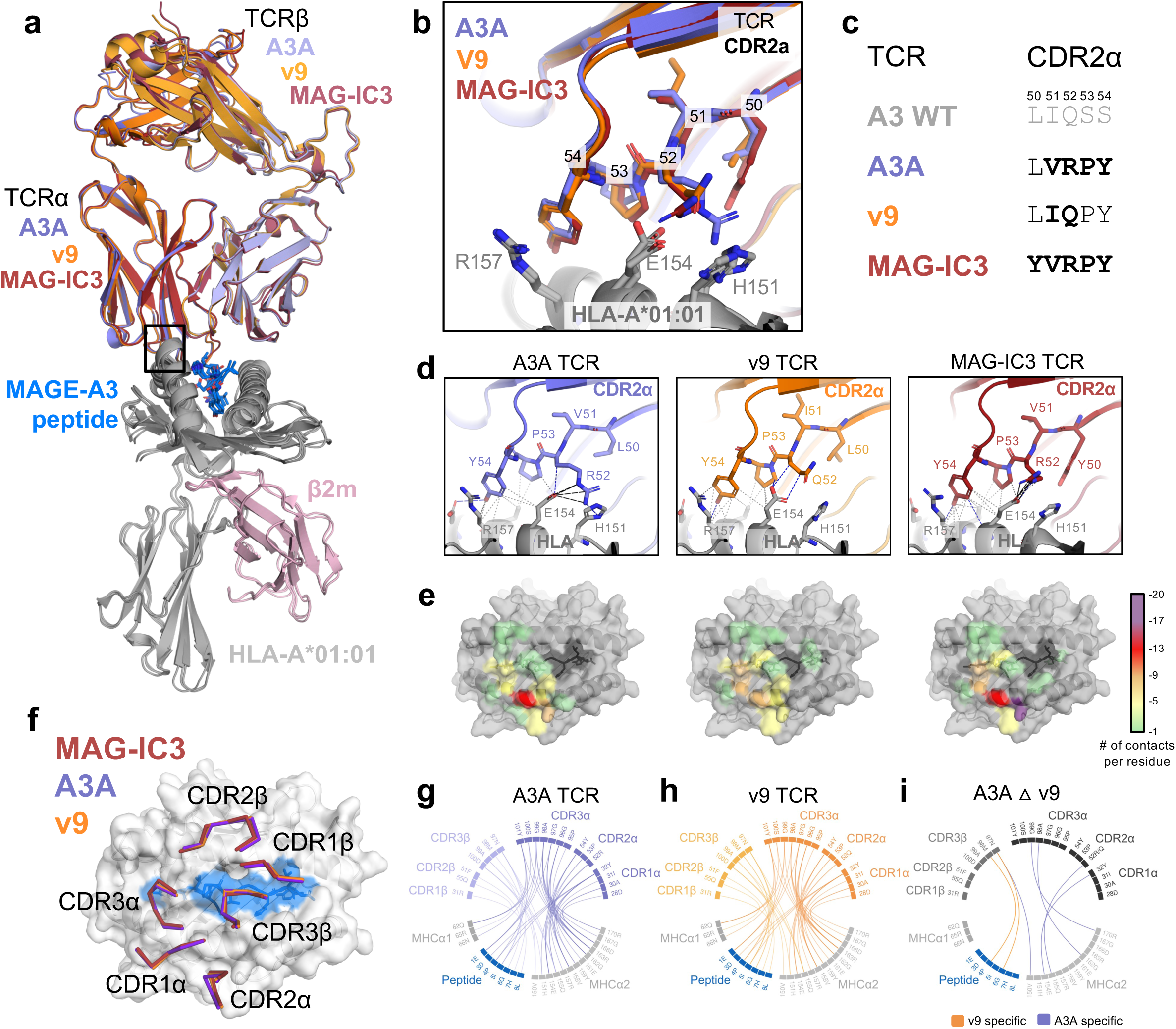
Structural comparison of the A3A and v9 TCR–MAGE-A3–HLA-A*01:01 complexes. **a**, Superposition of the MAG-IC3 (PDB ID 5BRZ), A3A and v9 TCR-MAGE-A3-HLA-A*01:01 complex structures. The peptide is depicted in blue, β2m in pink, HLA-A*01:01 in grey and TCRs in purple (A3A), orange (v9) and red (MAG-IC3). **b,** Close-up view of the TCR/CDR2α- HLA-A*01:01 interface, corresponding the to the black rectangle in **(a)**. **c,** Sequence alignment of the CDR2α residues in the A3 WT, A3A, v9 and MAG-IC3 TCRs. Highlighted in bold are the residues that are different to the A3 WT sequence. **d,** Close-up views of the TCR/CDR2α- HLA-A*01:01 interfaces, highlighting the residues that form interactions with the HLA. The A3A TCR is depicted in purple (left panel), v9 TCR in orange (central panel) and MAG-IC3 TCR in red (right panel). Hydrogen bonds, salt bridges, van-der-Waals and hydrophobic interactions are shown as dashed lines. **e,** Surface representation of the MAGE-A3-HLA-A*01:01 complex, shown in grey (HLA-A*01:01) and dark grey (MAGE-A3 peptide). The interaction footprints of the respective TCRs (A3A, left panel; v9, central panel; and MAG-IC3, right panel) are shown in colour. Interacting residues on the MAGE-A3-HLA-A*01:01 surface are coloured depending on how many contacts are formed with the TCR. No contacts (grey), 1-4 (green), 5-8 (yellow), 9-12 (orange), 13-16 (red), 16-20 (purple). **f,** Top views of CDRs from the three TCR/MAGE-A3-HLA-A*01:01 complexes (MAG-IC3 red; A3A purple; v9 orange). **g-i,** Chord diagrams representing the molecular interactions for **(g)** A3A and **(h)** v9, and **(i)** shows the symmetric difference between A3A and v9.

In the case of A3A, the engineered CDR2α sequence formed extensive interactions with the MHC α2-helix: the side chain of Arg52 extended towards the MHC and formed 2 Hydrogen bonds, 4 salt bridges and 3 polar interactions with residues His151 and Glu154 (**Fig. 4d; Supplementary Table 2**), while residues Pro53 and Tyr54 contributed further contacts to the MHC α2-helix. Reverse engineering A3A to v9 (LVRPY -> LIQPY) resulted in the loss of 3 Hydrogen bonds and 4 salt bridges between the CDR2α and the MHC α2-helix, while the contacts provided by Pro53 and Tyr54 were maintained (**Fig. 4d**). These additional interactions, which would largely be absent in the A3 WT complex (LIQSS), suggest an explanation for the differences in binding affinities of A3A, v9, and A3 WT for MAGE-A3 (**Extended Data Fig. 8a,b,d**; K_D_ 1.52 μM vs 23.87 vs >500 μM^7^).

A3A and v9 provide two highly similar solutions to MAGE-A3 recognition that nevertheless resulted in distinct Titin cross-reactivity. To add additional context to the impact of the subtle changes at the TCR/pMHC interface, we compared the pMHC footprints of v9 and A3A to their original wild-type sequence, A3 WT, as well as an affinity-tuned version of the A3A, named MAG-IC3^22^ (CDR2α sequence: YVRPY) that recognises both MAGE-A3 and Titin with approximately 1,000-fold higher affinity than A3A (7.1 and 76.7 nM K_D_, respectively; PDB ID 5BRZ). Since there are no available structures of the low-affinity A3 WT, we generated a prediction of the A3 WT complex with MAGE-A3-HLA-A*01:01 using AlphaFold3 (AF3; **Extended Data Fig. 10a-c**). We found that A3 WT (LIQSS) displayed a more balanced footprint compared to A3A and MAG-IC3, with proportionally fewer interactions engaging the MHC (72% for A3 WT vs >80% for A3A and MAG-IC3; **Extended Data Fig. 10c-e; Fig. 4e**). In contrast, v9 re-balanced the ratio of MHC to peptide contacts (73% of contacts with MHC; **Fig. 4e; Extended Data Fig. 10d-l**). Taken together, the affinity data and contact interface analysis suggested the gains in affinity obtained with the engineering of A3A and MAG-IC3 resulted from interactions mediated with MHC residues Arg52, Pro53 and Tyr54. However, these newly established contacts also shifted the balance of the interaction away from peptide-specific contacts. In contrast, v9 maintained recognition of the MHC via Pro53 and Tyr54 but lost the aforementioned contacts mediated by Arg52, likely contributing to a reduction in affinity but making the interaction more peptide-dependent.

Finally, we mapped all interactions between the CDR loops of the A3A and v9 TCRs and the MAGE-A3-HLA-A*01:01 complex (**Fig. 4f-h**). The majority of the CDR contacts were conserved between the two structures, but several notable differences emerged. Unexpectedly, most of these unique contacts arose outside the CDR2α loop (**Fig. 4i**). v9 CDR3β formed two additional contacts with the peptide, whereas A3A CDR3α and CDR1α formed two and one additional contacts with the MHC, respectively. Most strikingly, CDR3β Asn97, a residue shared by the two TCRs, engaged the peptide in the v9 complex but redirected to the MHC in the A3A complex, illustrating that the same residue in the CDR3β loop was influenced by the reversed residues in the CDR2α. Together, these contact differences suggested that the CDR2α substitutions were not only acting locally but were also propagating across the TCR, reshaping how distal CDR loops engaged the pMHC surface.

### Molecular dynamics simulations of A3A and v9 TCR/MAGE-A3-HLA-A*01:01 interactions

To better understand these distinct interaction networks, we used all-atom molecular dynamics (MD) simulations to interrogate the dynamics of the forces that shape the contact interface, and specifically, whether the mutated residues in CDR2α related to differences in conformational states across all CDRs. Prior to conformational analysis, RMSD and RMSF analyses confirmed that all simulations were stable and that the CDR2α mutations did not introduce differences in backbone flexibility in the mutated region (**Extended Data Fig. 11a,b**). We next applied time-lagged independent component analysis (tICA) and Markov State Models (MSM) to the conformational ensembles of the A3A and v9 TCRs in the unbound and bound states to MAGE-A3/HLA-A*01:01. In line with the crystallographic data, tICA-MSM conformational dynamics analysis revealed that the CDR2α backbone conformation was unchanged in v9 relative to A3A despite the differences in residues 51 and 52 (**Fig. 5a,b**). Instead, the CDR2α mutations in v9 propagated across the TCR, resulting in new conformational states for both CDR3α (**Fig. 5c,d**) and CDR3β (**Fig. 5e,f**).

**Fig. 5.**
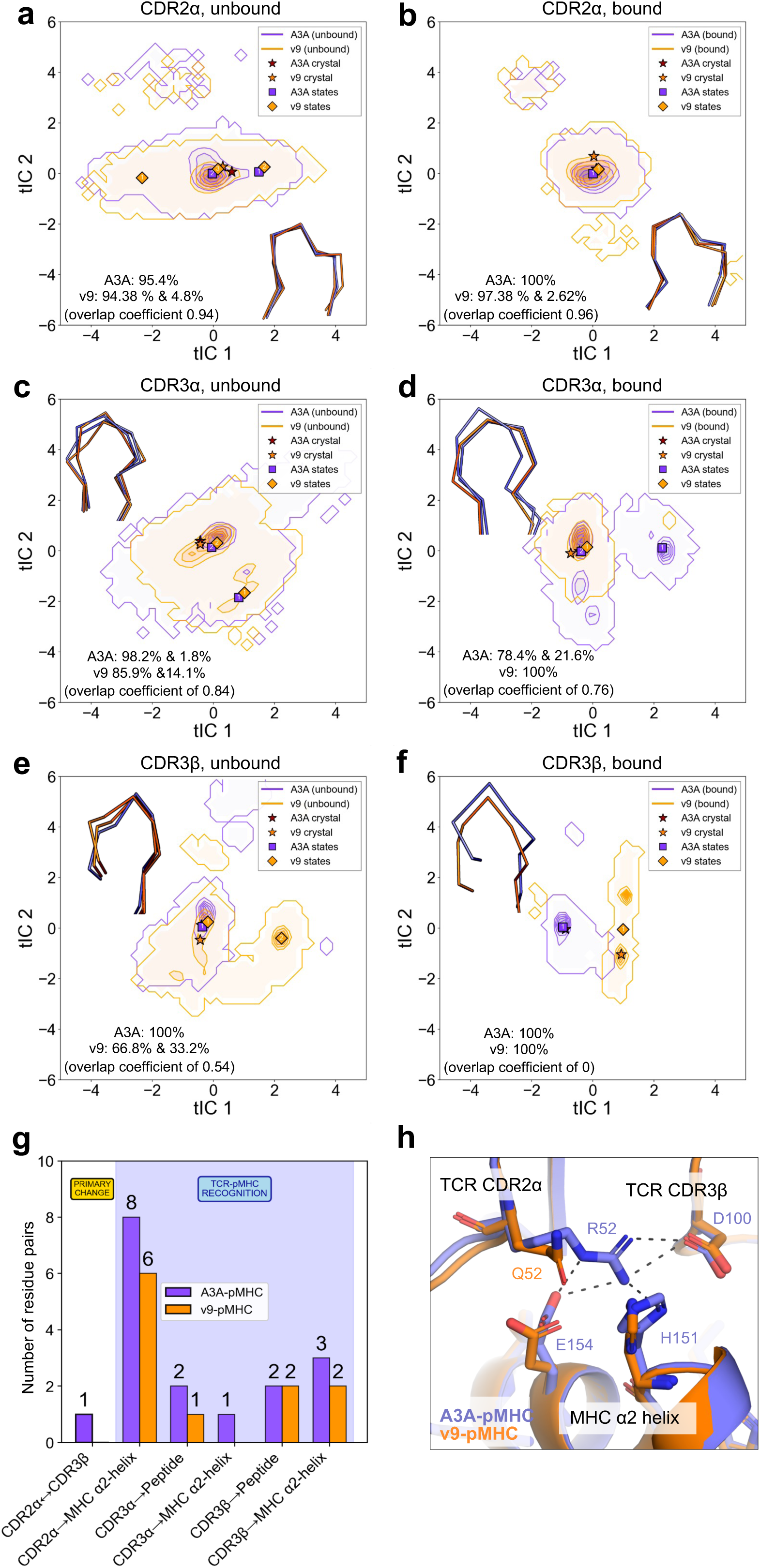
Time-lagged independent component (tIC) analysis of CDR loop conformational dynamics, contact network comparison, and structural representation in A3A and v9 unbound and bound TCRs. **a-f**, Contour density plots showing conformational distributions for A3A (purple) and v9 (orange). Left panel unbound, right panel bound. Stars represent crystal structures, diamonds represent simulation-identified centroid states. The numbers in each panel report the population percentage of each identified conformational macrostate and Bhattacharyya overlap coefficient between A3A and v9 distributions. **(a,b)** Conformational distributions for the CDR2α loop in A3A and v9 in **(a)** unbound and **(b)** bound states. **(c,d)** Conformational distributions for the CDR3α loop in **(c)** unbound and **(d)** bound states. **(e,f)** Conformational distributions for the CDR3β loop in **(e)** unbound and **(f)** bound states. **g,** Contact network comparison: bar chart showing the number of residue pair interactions (≥25% persistence, 4.5Å cutoff) between A3A/pMHC (purple) and v9/pMHC (orange). The bars represent different contact categories: primary change (CDR2α↔CDR3β) and TCR-pMHC recognition contacts (CDR loops to pMHC). **h,** Structural representation of the A3A (purple) and v9 (orange) CDR loops showing CDR2α and CDR3β regions. Grey dashed lines indicate the hydrogen bonding interaction between CDR2α- Arg52 and CDR3β-Asp100 present in the A3A/pMHC structure.

For the CDR3α loop, the conformational ensembles of A3A and v9 were similar in the unbound state (overlap coefficient 0.84, **Fig. 5c**). Upon binding to pMHC, the A3A CDR3α developed two-state behaviour whereas the v9 CDR3α remained in a single state (**Fig. 5d**), and the conformational overlap between the TCRs decreased from 0.84 to 0.76 (**Fig. 5c,d**). The difference in dynamics was even more pronounced for the CDR3β loop: in the unbound state, v9 adopted distinct two-state behaviour whereas A3A remained in a single state (overlap coefficient of 0.54, **Fig. 5e**). Upon binding, both A3A and v9 exhibited stable single-state behaviours, but their conformations showed no overlap, indicative of distinct bound conformations despite having the same CDR3β sequence (**Fig. 5f**). In contrast, CDR1α, CDR1β and CDR2β showed largely consistent conformational dynamics between A3A and v9 in both the unbound and bound states (**Extended Data Fig. 12a-f**). Chapman-Kolmogorov tests confirmed that MSM-predicted transition probabilities were in close agreement with direct MD estimates across multiple lag times for all CDR loops in both the unbound and bound states, validating the Markovian assumption underlying each model (**Extended Data Fig. 13, Extended Data Fig. 14**).

These time-course analyses confirmed that the changes introduced during the reverse engineering of the CDR2α loop were propagated throughout the TCR and remained stable over long MD time courses. However, in isolation, these analyses did not account for the altered patterns of cross-reactivity. To investigate this, we quantified persistent residue-pair interactions (defined as within a 4.5 Å cutoff, ≥25% persistence, and side-chain only interactions) over the entire simulation and divided them into TCR/pMHC recognition contacts (CDR-pMHC) or intra-TCR contacts (CDR-CDR and CDR-framework) (**Extended Data Fig. 12g**, **Supplementary Table 3**). From this analysis, the single most consequential difference between A3A and v9 was the loss of an intra-TCR side-chain contact between CDR2α Arg52 and CDR3β Asp100 (**Fig. 5g,h**). Overall, v9 CDRs demonstrated fewer stable contacts with both the peptide and MHC than A3A (**Fig. 5g**), with CDR2α, CDR3α, and CDR3β collectively forming fewer contacts across both targets (16 for A3A vs 11 for v9). The reduction was driven primarily by the loss of the intra-TCR CDR2α Arg52-CDR3β Asp100 contact in v9, as well as by fewer CDR2α contacts with the MHC α2-helix (8 vs 6), one fewer CDR3α-peptide contact (2 vs 1) resulting in the loss of the Tyr101-Pro4 contact, a complete loss of CDR3α-MHC α2-helix contacts in v9 (1 vs 0), and one fewer CDR3β/MHC α2-helix contact (3 vs 2) (**Fig. 5g, Extended Data Fig. 15a-e**). Whilst the total number of contacts formed by CDR3β with the peptide were identical between the two TCRs (2 each; **Fig. 5g**), the chemical composition of the interface was altered, as the Asn97-His7 contact present in A3A was absent in v9 and replaced by a novel Met98-Pro4 (**Extended Data Fig. 15f**). Notably, Pro4 was recognised by both TCRs but via different residues and CDRs: Tyr101 CDR3α in A3A and Met98 CDR3β in v9, suggesting a clear shift in how v9 engaged this peptide position rather than a complete loss of interaction (**Extended Data Fig. 15c-f**). Taken together, this modified interaction network, co-occurring with disruption of the intra-TCR CDR2α–CDR3β contact, is consistent with allosteric coupling between the CDR2α and CDR3 loops and provides a structural rationale for v9’s rebalanced specificity profile relative to A3A.

## Discussion

The affinity-enhanced A3A TCR differs from its wild-type precursor by only four residues in the CDR2α, yet this engineering step was sufficient to compromise cross-reactivity and to introduce recognition of multiple wild-type peptides, including a Titin-derived peptide that led to fatal cardiotoxicity^7^. Rather than viewing such affinity-enhanced receptors as failed endpoints, we demonstrate that they can instead serve as starting points for reverse engineering. By systematically reverting engineered residues to wild-type, we reconstructed a high-resolution cross-reactivity map that links the low-affinity A3 WT to its affinity-enhanced A3A. To our knowledge, this represents one of the most comprehensive specificity maps of closely related TCR/pMHC interactions, integrating yeast-display profiling, functional validation, structural analysis, and molecular dynamics. This approach offers a blueprint to rescue engineered TCRs whose mutational distance from a thymically-selected precursor is small enough to be exhaustively explored, complementing *de novo* discovery, library-based functional screening^25^, and biophysical engineering approaches such as catch-bond tuning^28^.

Large-scale yeast-display screening, combined with quantitative motif analysis and functional validation^9,10,27^, enabled prioritisation of variants based on both peptide degeneracy and functional reactivity. This allowed rational selection of the v9 variant, which retained potent MAGE-A3 recognition while eliminating reactivity to Titin and other off-target self-antigens. Across the intermediate TCR variants, two distinct molecular solutions emerged that preserved robust MAGE reactivity while minimising cross-reactivity. The first combined restoration of the wild-type Gln52 but maintained Pro53 and Tyr54 (v9 LIQPY); the second retained the engineered Arg52 but restored at least one of the original Ser53 or Ser54 (v3 LIRSS, v7 LIRSY, v8 LIRPS). That two chemically distinct CDR2α configurations converge on similar specificity profiles suggests that the engineered A3A occupies a local optimum for affinity but not for specificity, and that more than one return path to a safer receptor exists within a four-residue Hamming distance. These minimal sequence permutations were sufficient to reshape peptide degeneracy profiles, demonstrating that subtle modulation of the CDR2α loop can rebalance sensitivity and specificity without altering overall docking geometry. Such side-by-side specificity mapping provides a scalable method for benchmarking engineered TCRs and prioritising candidates for further clinical development.

Mechanistically, our structural and MD analyses reveal that cross-reactivity is not dictated solely by direct peptide contacts. Despite near-identical docking geometries, subtle CDR2α mutations altered the stability of MHC α2-helix contacts^29^ and disrupted intra-CDR contacts between CDR2α and CDR3β. The accompanying reshaping of CDR3 conformational ensembles, observed across simulation timescales, is consistent with allosteric coupling between CDR2α and the CDR3 loops. CDR2-mediated effects on CDR3 dynamics have been described in other TCR systems^15,16^, and our data extend this principle by identifying a specific intra-TCR contact whose disruption tracks with reprogrammed peptide specificity.

Our results also inform broader TCR engineering strategies^2^. Affinity maturation has been widely pursued to enhance therapeutic potency, yet supraphysiological affinity often introduces increased off-target recognition^5,6^. Notably, although v9 displayed an approximately 10-fold lower affinity for MAGE-A3 by SPR compared with A3A, this reduction did not compromise cytotoxicity; v9-mediated killing of peptide-pulsed and endogenously expressing targets was comparable to A3A across multiple effector-to-target ratios. These findings reinforce the concept that supraphysiological affinity is not required for effective tumour cell killing^14,30^. Instead, tuning interaction stability and dynamic coupling may be sufficient to achieve therapeutic potency while reducing off-target risk. The kinetic dissociation profiles (particularly when considered in 2D-systems), catch-bond behaviour, and CD8 co-receptor coupling may all contribute, and dissecting these will be important for translating reverse-engineering into a predictive design framework.

Importantly, while v9 eliminated Titin reactivity and most A3A-acquired cross-reactivities, it retained the modest reactivity of the A3 WT toward antigens such as hFat2 and PLD5^12^. This observation underscores a critical principle at the core of this approach, and an important limitation: engineering can remove newly introduced cross-reactivity, but it does not erase the intrinsic degeneracy of the wild-type receptor. Comprehensive mapping of native TCR cross-reactivity should therefore precede affinity enhancement to avoid propagating or amplifying pre-existing liabilities.

Several limitations should be noted. The A3 WT complex structure was predicted by AlphaFold3 rather than determined experimentally. The allosteric mechanism is inferred from static structures and MD simulations but has not been tested by mutagenesis of the CDR2α-CDR3β contact. Although v9 was tested against 12 confirmed A3A agonists and the top 55 v9-predicted candidates, an exhaustive genome-wide screen was not performed (e.g. T-Scan^12^ or TCR-MAP^13^). Finally, although residual low-level reactivity to endogenous antigens cannot be fully excluded without altering the overall docking geometry, reducing the interaction’s overall stability may be sufficient to mitigate off-target toxicity. Future preclinical models will be required to define these thresholds. More broadly, our study provides a paradigm for dissecting and refining TCR specificity through stepwise reverse engineering, advancing both mechanistic understanding and the rational development of safer TCR-based immunotherapies.

---

**Extended Data Fig. 1.**
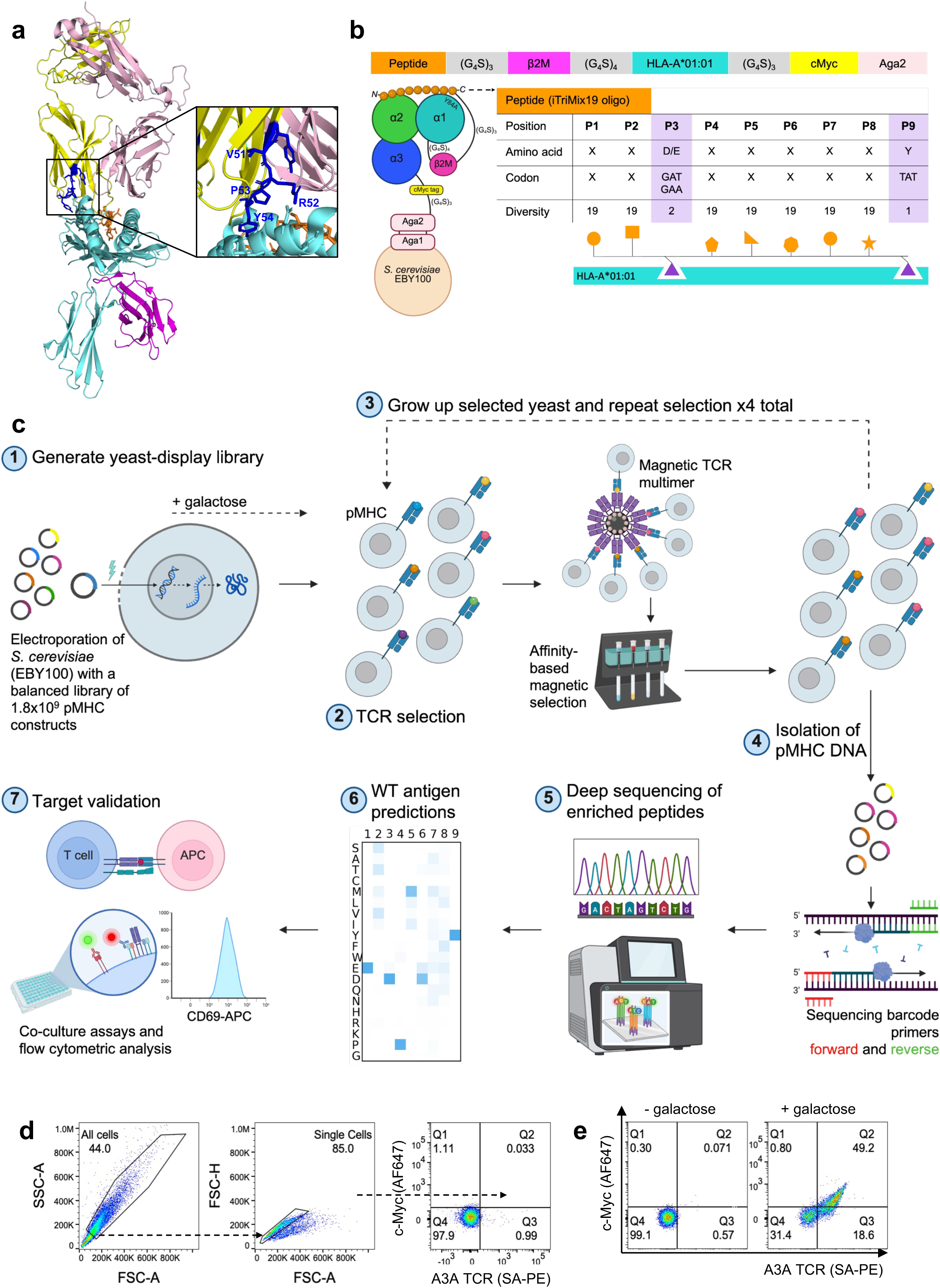
Development of a 9-mer-HLA-A*01:01 yeast-display library. **a**, Cartoon depiction of an A3A-like TCR in complex with MAGE-A3-HLA-A*01:01 (PDB: 5BRZ, MAG-IC3). TCRα chain in yellow, TCRβ chain in light pink, MAGE-A3 peptide in orange, HLA-A*01:01 in aquamarine, β2M in magenta. The CDR2α loop is in dark blue with residues shared with A3A annotated (hence R50 is not annotated: A3A, Leu50, LVRPY; MAG-IC3, Tyr50, YVRPY). **b,** Top: combinatorial 9-mer- A*01:01 library expression construct. Bottom: schematic of the 9-mer-A*01:01 combinatorial library, demonstrating how a diversity of 1.8x10^9^ 9-mer peptides was achieved using iTriMix19 oligos: one codon per amino acid (excluding cysteines) was randomly allocated to each position in the peptide, excluding the anchors P3 and P9, amounting to 1.8 × 10^9^ peptides (19^7^ × 2 × 1). Anchors were fixed based on the known peptide preferences of HLA-A*01:01: D = Glu, E = Asp, Y= Tyr. **c,** Schematic depiction of the pMHC yeast-display method: (1) EBY100 yeast cells were electroporated with the iTriMix19 library of 1.8x10^9^ 9-mer-A*01:01 constructs. (2) Affinity-based magnetic selection with magnetic bead-multimerised TCRs were used to enrich yeast cells displaying agonist peptides. (3) Step 2 was repeated three more times, further enriching for yeast cells displaying agonist peptides. (4) The 9-mer-A*01:01 plasmids were isolated from the yeast cells enriched per round, and unique sequencing barcodes were added by PCR. (5) The isolated, barcoded 9-mer-A*01:01 constructs were deep sequenced for identification of the enriched peptides. (6) *In silico* motif similarity analysis was applied to the deep sequencing reads to predict the corresponding wild-type antigens. (7) Co-culture activation assays were conducted to validate the predicted wild-type antigens. **d,** General flow cytometry gating strategy for non-induced yeast transformed with the MAGE-A3-HLA-A*01:01 scaffold (stained with anti-c-Myc-AF647 and 1 μM SA-PE A3A-tetramer). **e,** Flow cytometry dot plots showing binding of the A3A-tetramer to induced yeast expressing MAGE-A3- A*01:01 complexes, detected by c-Myc expression (stained with anti-c-Myc-AF647 and 1 μM SA-PE A3A-tetramer).

**Extended Data Fig. 2.**
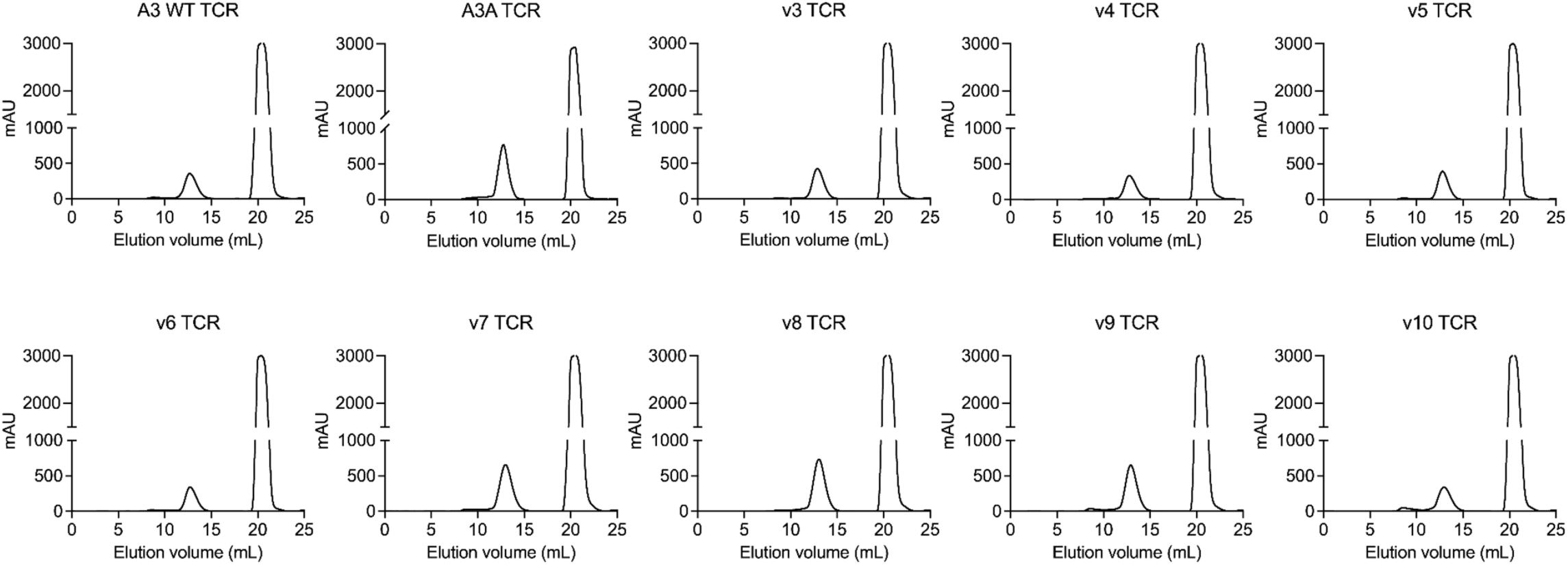
Production of recombinant A3A-variant TCRs. Size-exclusion chromatography profiles of the A3A-variant TCRs following expression (in the Expi293 mammalian system), purification, and biotinylation. The left peak eluting at approx. 13 mL corresponds to the respective TCRs, and the right peak eluting after 20 mL corresponds to free biotin. These recombinant TCRs were used in the pMHC yeast-display screens.

**Extended Data Fig. 3.**
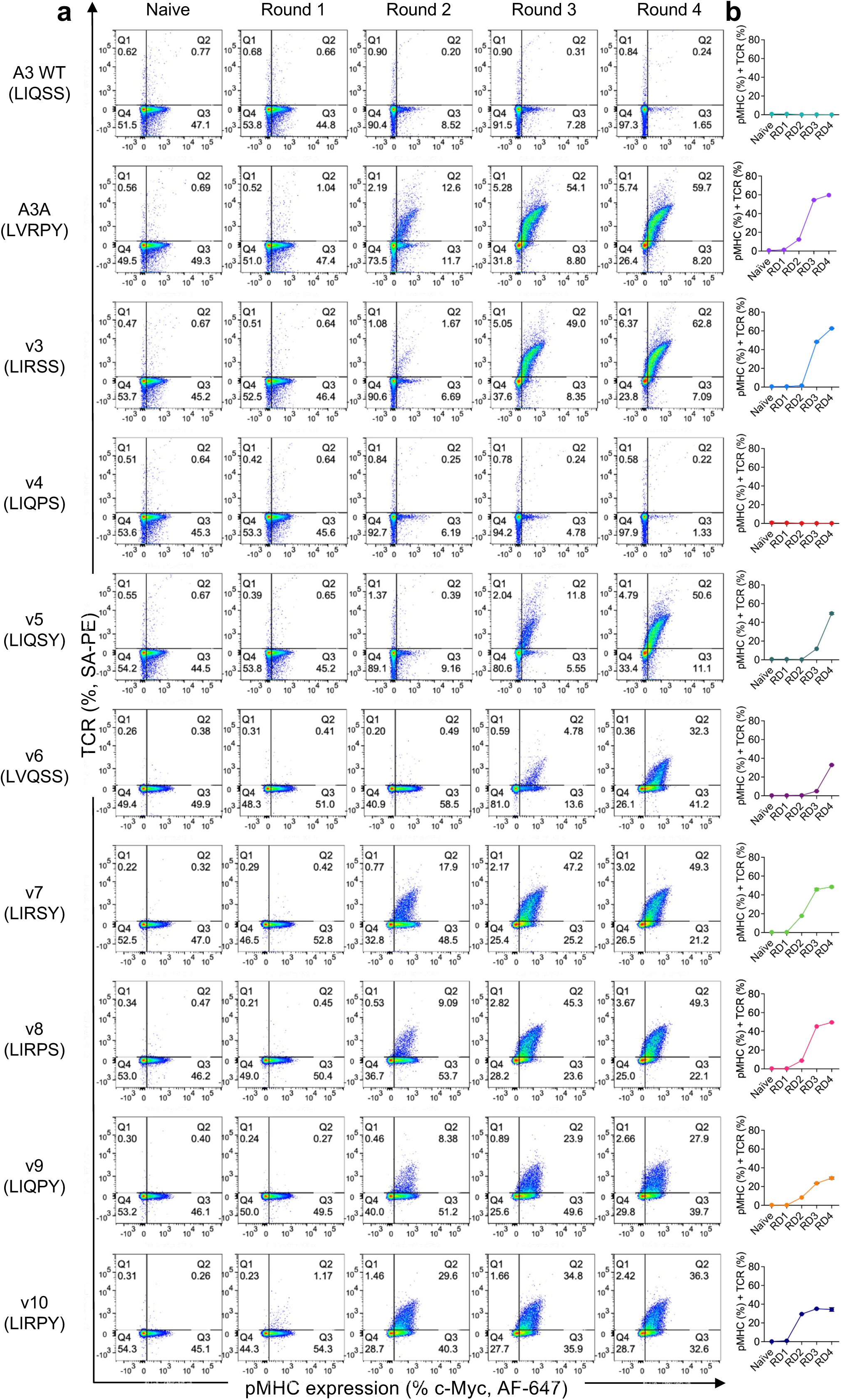
Tetramer staining of all 10 A3A-variant pMHC yeast-display screens. **a**, Flow cytometry plots depicting total A3A TCR tetramer staining of naïve yeast (the original 9-mer-A*01:01 library prior to TCR selection) and after all four selection rounds (R1-R4) with the ten A3A TCR variants. Yeast were stained with anti-c-Myc-AF647 and 1 μM SA-PE TCR-tetramer. **b,** Graphical depictions of the flow cytometry plots showing the percentage of yeast displaying TCR-tetramer-stained pMHC molecules at each selection round (i.e. the double positive populations). Colours match with Fig. 1c.

**Extended Data Fig. 4.**
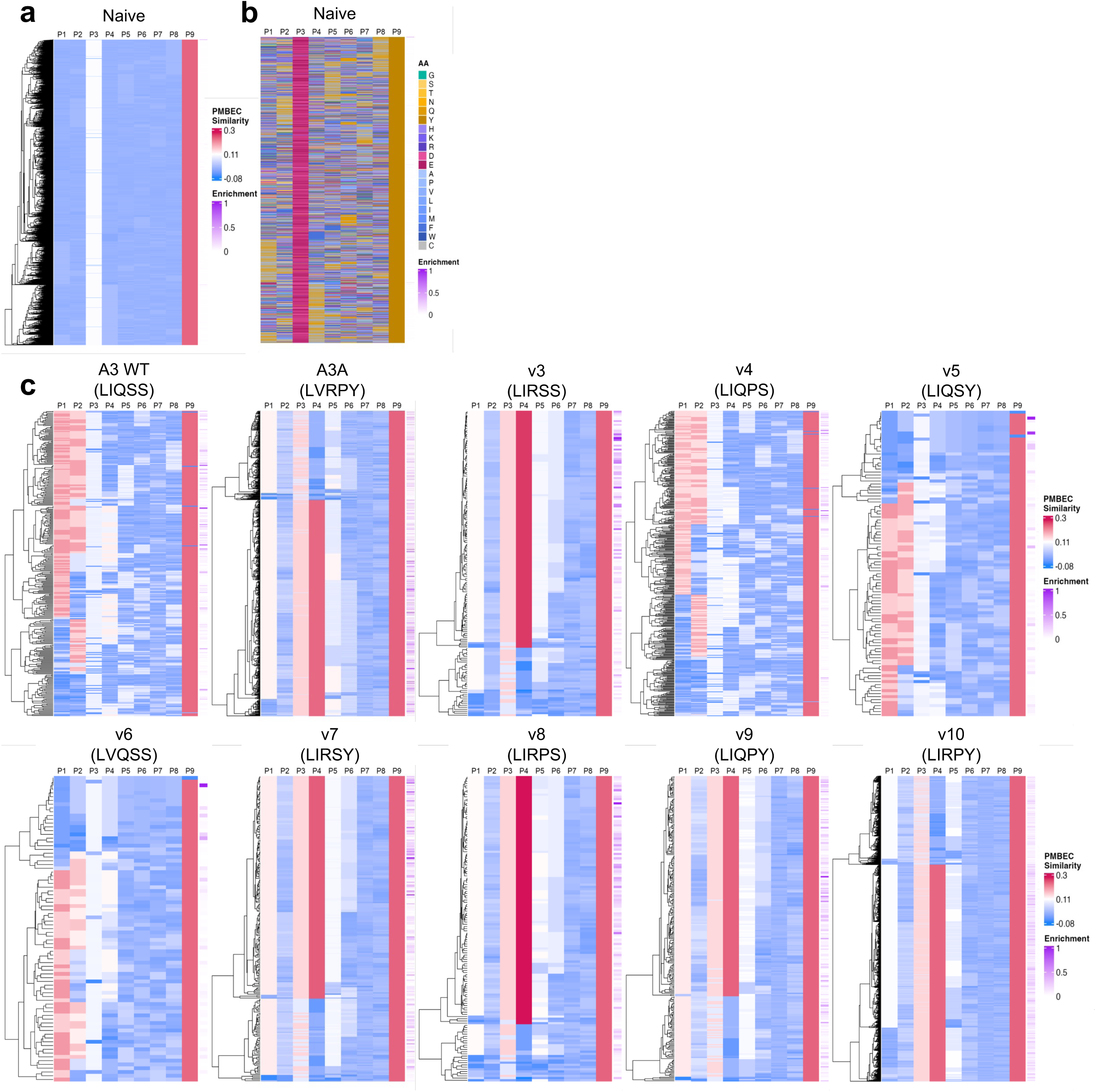
Chemical similarity heatmaps displaying the pMHC yeast-display deep sequencing data. **a-b**, Heatmaps displaying the top 5000 peptides from the naive 9-mer-A*01:01 yeast-display library. **a,** The PMBEC score for each individual peptide against all other peptides was calculated, the average was taken, and the peptides were grouped using hierarchical clustering. **b,** Each amino acid was then assigned a different colour (grouped by chemical similarity), as displayed by the “AA” key, with larger amino acids assigned darker colours. The single-letter abbreviations for the amino acid residues are as follows: A, Ala; C, Cys; D, Asp; E, Glu; F, Phe; G, Gly; H, His; I, Ile; K, Lys; L, Leu; M, Met; N, Asn; P, Pro; Q, Gln; R, Arg; S, Ser; T, Thr; V, Val; W, Trp; and Y, Tyr. The enrichment level of each peptide is shown on the far right of each heatmap, with darker purple representing a higher frequency of that peptide out of the top 95% enriched peptides. **c,** Chemical similarity heatmaps comparing the top 95% of enriched peptides at round 4 from the pMHC yeast-display deep sequencing data from all ten TCR screens. The PMBEC similarity score for each peptide position was calculated by taking an enrichment-normalised average of all peptides, which is shown in Fig. 2b.

**Extended Data Fig. 5.**
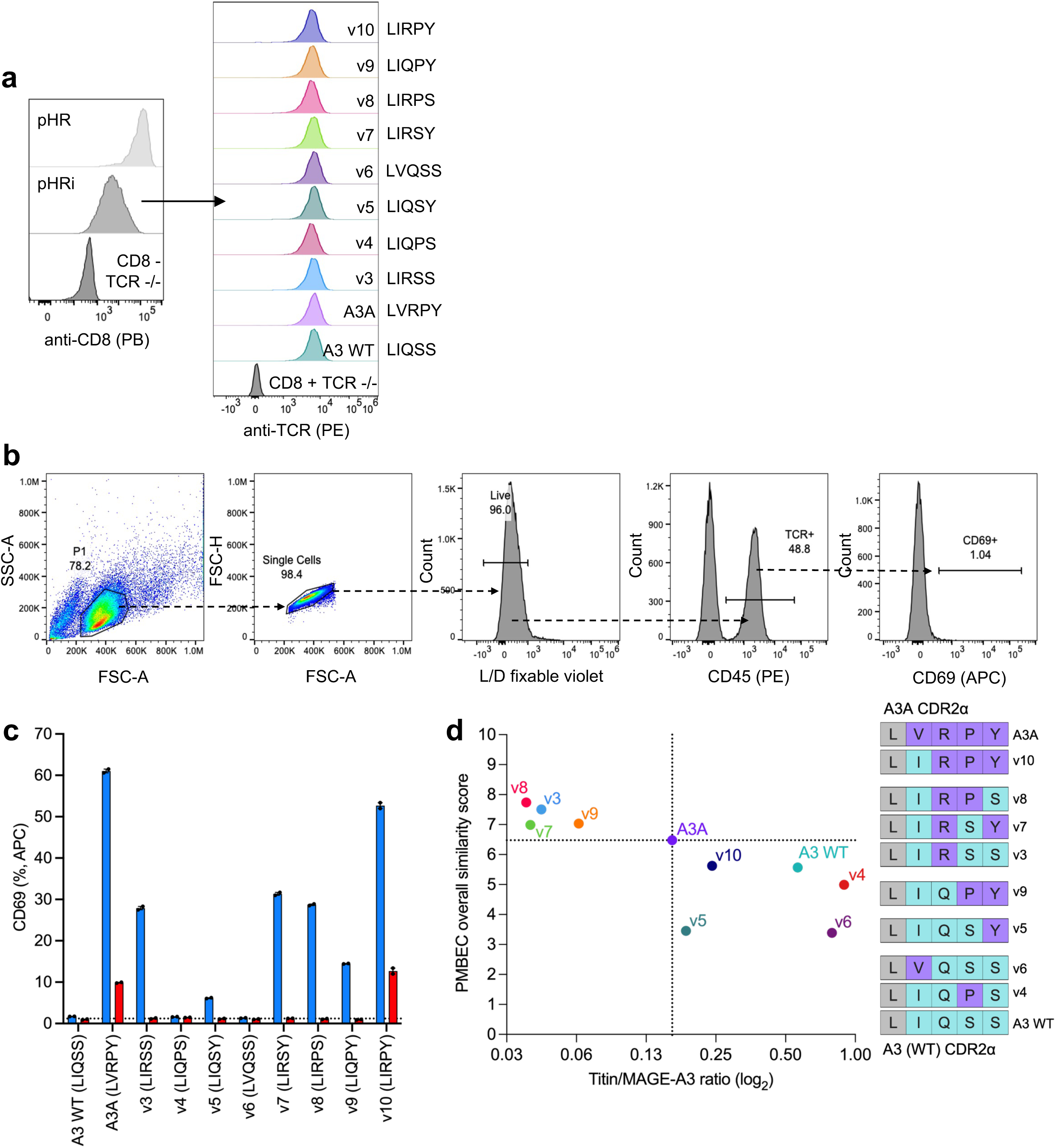
Transduction of cell lines for co-culture assays. **a**, The left panel shows the expression levels of CD8 on the surface of TCR-/- Jurkats transduced with the lentiviral pHR vector (SFFV promoter) or pHRi inducible vector (ecdysone-inducible promoter). In the absence of ecdysone the pHRi vector is transcribed at low levels, achieving physiological CD8 expression levels. These pHRi-transduced cells were then transduced with the ten TCR variants (A3 WT, A3A, and v3-v10), and the right panel shows the expression levels of these respective TCRs. **b,** Gating strategy for the co-culture assays, where peptide-pulsed A1-K562s were used to stimulate A3A-variant-Jurkat T cells at an E/T ratio of 1:1. The cells were stained with Live/Dead fixable violet dye, anti-CD45-PE (to select the Jurkat T cells), and anti-CD69-APC (to detect Jurkat T cell activation). **c,** A1-K562s were pulsed with 100μM of the peptides MAGE-A3, Titin, or a control non-agonist peptide (dotted line) for 2h. The ten CD8+ A3A-variant-Jurkat cell lines were then added at an E/T ratio of 1:1 for 18h. Finally, anti-CD69 staining was performed on the ten A3A-variant-Jurkat T cell lines, enabling measurement of CD69 expression levels by flow cytometry. Error bars represent means of technical duplicates ± SEM, and the data are representative of three independent experiments. **d,** Co-culture activation data for each TCR (corresponding to the data in Fig. 2c) were colour-coded to the PMBEC similarity scores (Fig. 2b). For the co-culture assays, CD8+ TCR -/- Jurkat T cells were transduced with the ten TCR variants, and K562 antigen-presenting cells (APCs) were transduced with HLA-A*01:01 (A1-K562s). The A1-K562s were then pulsed with 100μM of the peptides MAGE-A3, Titin, or a control non-agonist peptide for 2h, and were used to stimulate the ten CD8+ A3A-variant- Jurkat T cell lines at an E/T ratio of 1:1 for 18h. Anti-CD69 surface expression was determined by flow cytometry.

**Extended Data Fig. 6.**
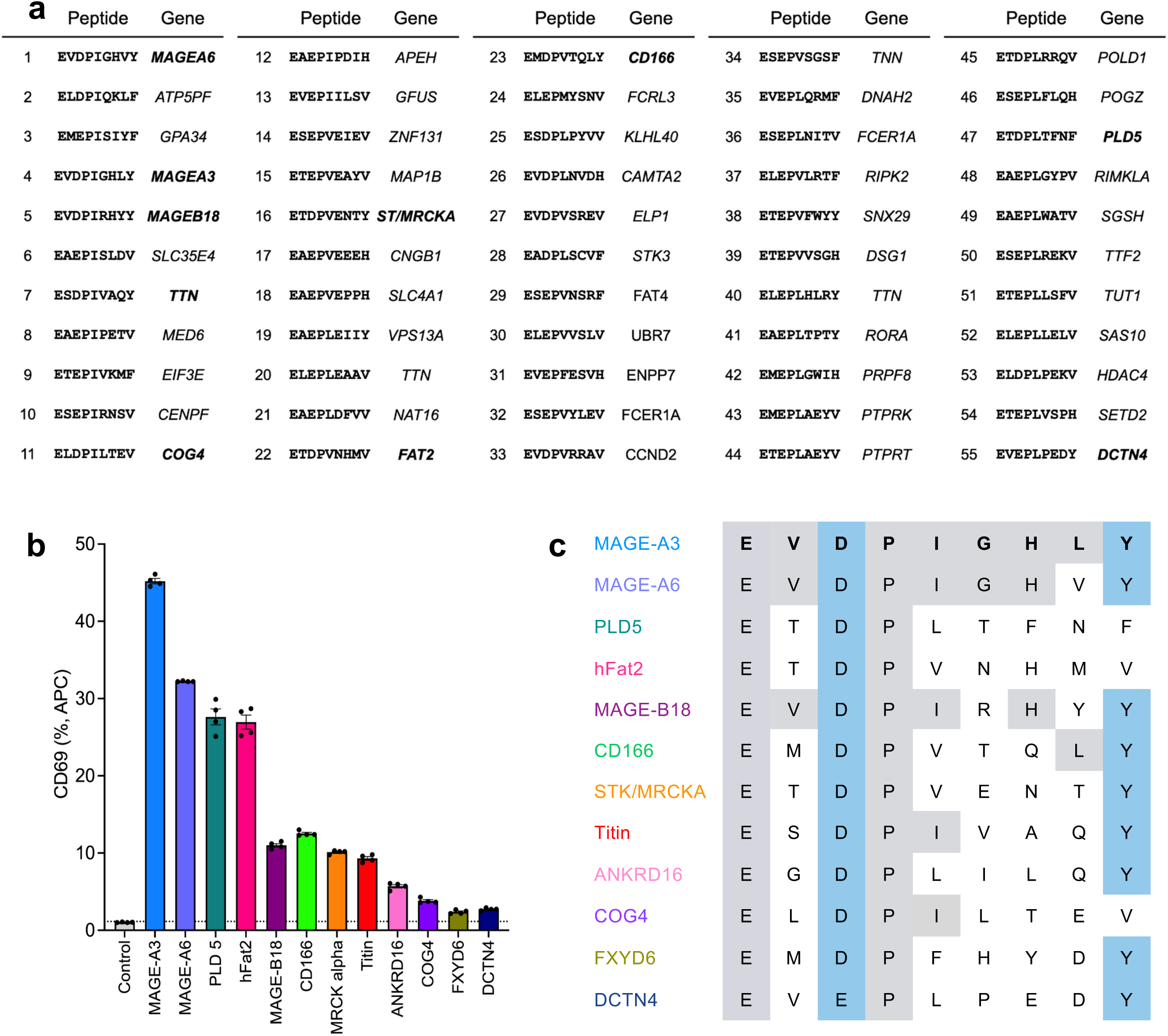
Validation of predicted A3A agonists from the 9-mer-A*01:01 yeast-display screen. **a**, The top-55 algorithm-predicted peptides from the human proteome, based on A3A’s pMHC yeast-display screen, were selected for subsequent validation in co-culture assays. The peptides are displayed in order of prediction ranking, with previously identified agonists of the A3A highlighted in bold, excluding no. 55 (*DCTN4*; Dynactin subunit 4) that was newly identified by our screen. **b,** A1-K562s were pulsed with 100μM of the 12 A3A-agonist peptides (agonists previously and newly identified by our yeast-display screen), or a control non-agonist peptide (grey bar, dotted line) for 2h. CD8+ A3A-variant-Jurkat cells were then added at an E/T ratio of 1:1 for 18h. Anti-CD69 staining was performed on the A3A-variant-Jurkat T cells, enabling measurement of CD69 expression levels by flow cytometry. Error bars represent means of technical replicates ± SEM pooled from two independent experiments. Statistical analysis by Welch’s ANOVA, with all peptides **p≤0.01. **c,** Amino acid sequences of the identified A3A TCR agonists in order of potency. Grey amino acids in common with the MAGE-A3 sequence, blue amino acids represent the fixed anchor residues at P3 and P9.

**Extended Data Fig. 7.**
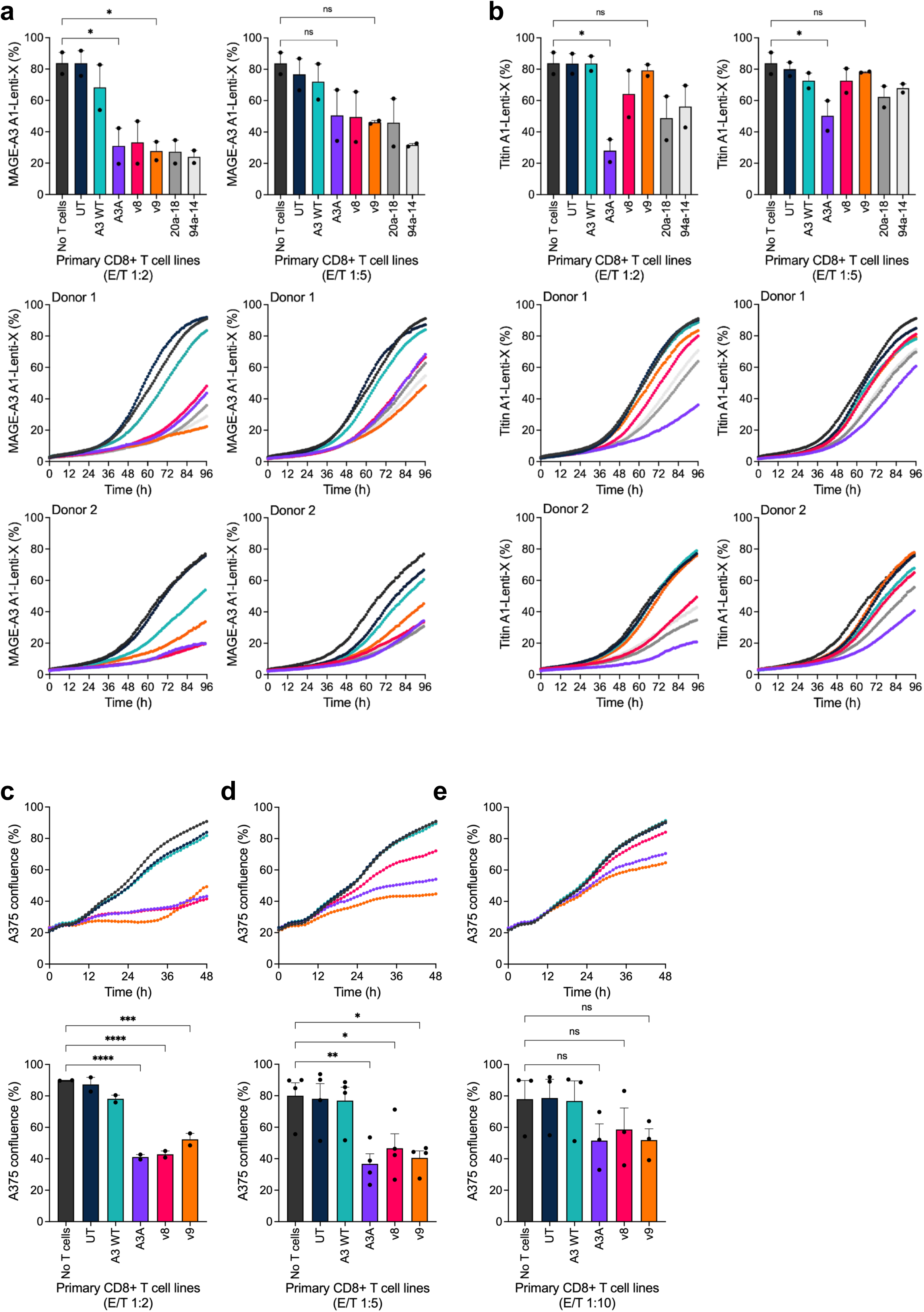
Cytotoxic capabilities of A3A-variant primary CD8+ T cell lines. Confluence of A1-Lenti-X cells pulsed with 50 μM of **a,** MAGE-A3 or **b,** Titin after 3 days cultured with or without A3A-variant primary CD8+ T cells from two healthy donors (top panel, data pooled from donors 1 and 2 showing % confluence at 96h; middle panel, healthy donor 1; bottom panel, healthy donor 2) at E/T ratios 1:2 (left column) or 1:5 (right column). UT = untransduced. Statistical analysis by ordinary oneway ANOVA, *p≤0.05, ns p>0.05. Error bars represent means ± SEM. Healthy donor 1 data and pooled E/T 1:2 data also shown in Fig. 3b-d. **c-e,** Top panel: confluence of A375 cells after 48h cultured with or without primary A3A-variant primary CD8+ T cells at E/T ratios **(c)** 1:2, **(d)** 1:5, and **(e)** 1:10. Bottom panel: data equivalent to top panel, pooled from two, four, and three independent experiments with different donors, respectively. Error bars represent means of biological replicates ± SEM. Statistical analysis by ordinary one-way ANOVA, ****p≤0.0001, ***p≤0.001, **p≤0.01, *p≤0.05, ns p>0.05.

**Extended Data Fig. 8.**
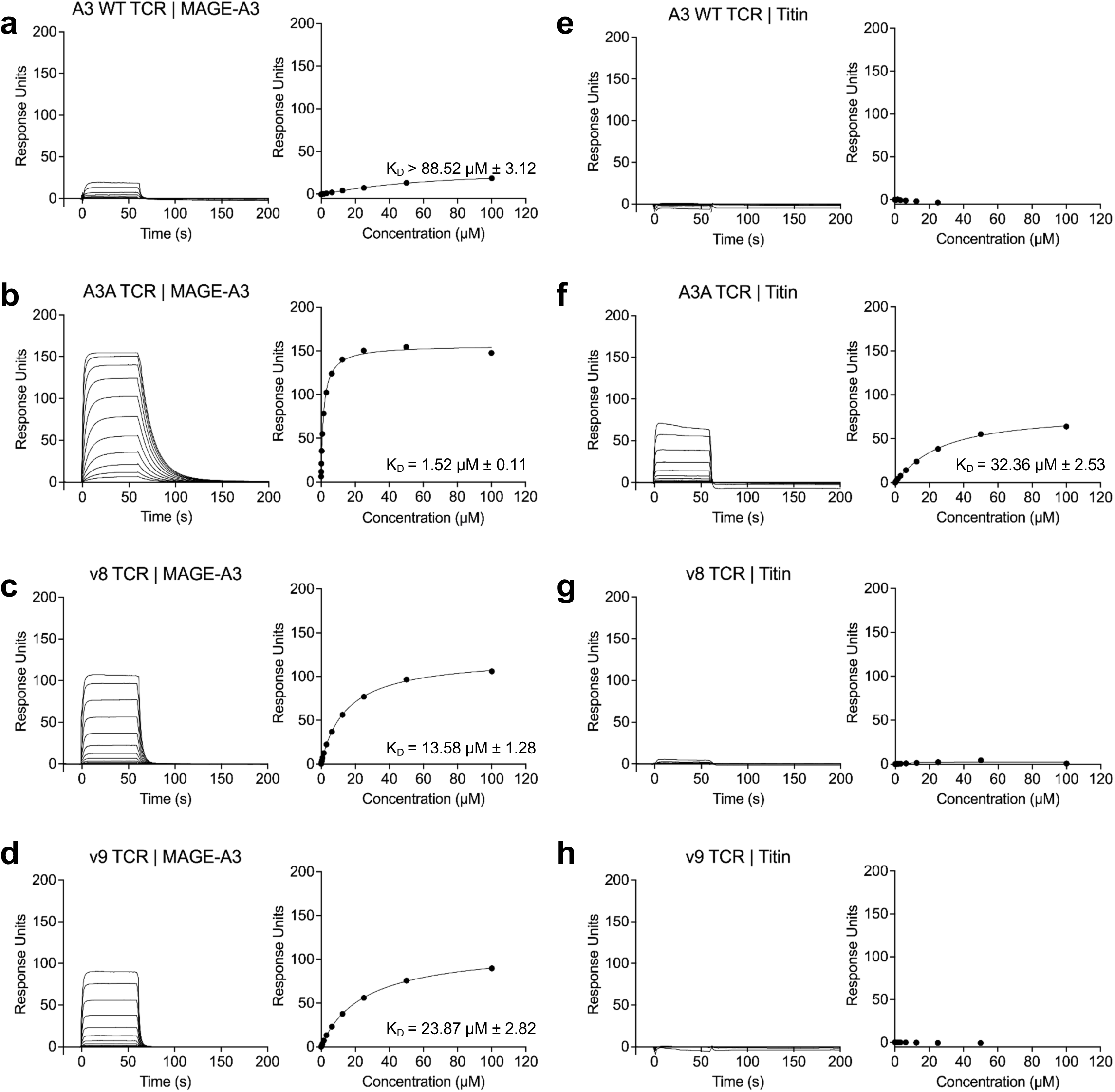
SPR sensorgrams of A3A-variant TCRs with MAGE-A3 and Titin-HLA-A*01:01 complexes. SPR binding curves of eight concentrations of A3 WT, A3A, v8 and v9 TCRs flown over **(a-d)** MAGE-A3-HLA-A*01:01 or **(e-h)** Titin-HLAA* 01:01 complexes, docked onto a Biacore CAP chip. Data are representative of three independent experiments, with Kds calculated as the mean ± SD of these three repeats.

**Extended Data Fig. 9.**
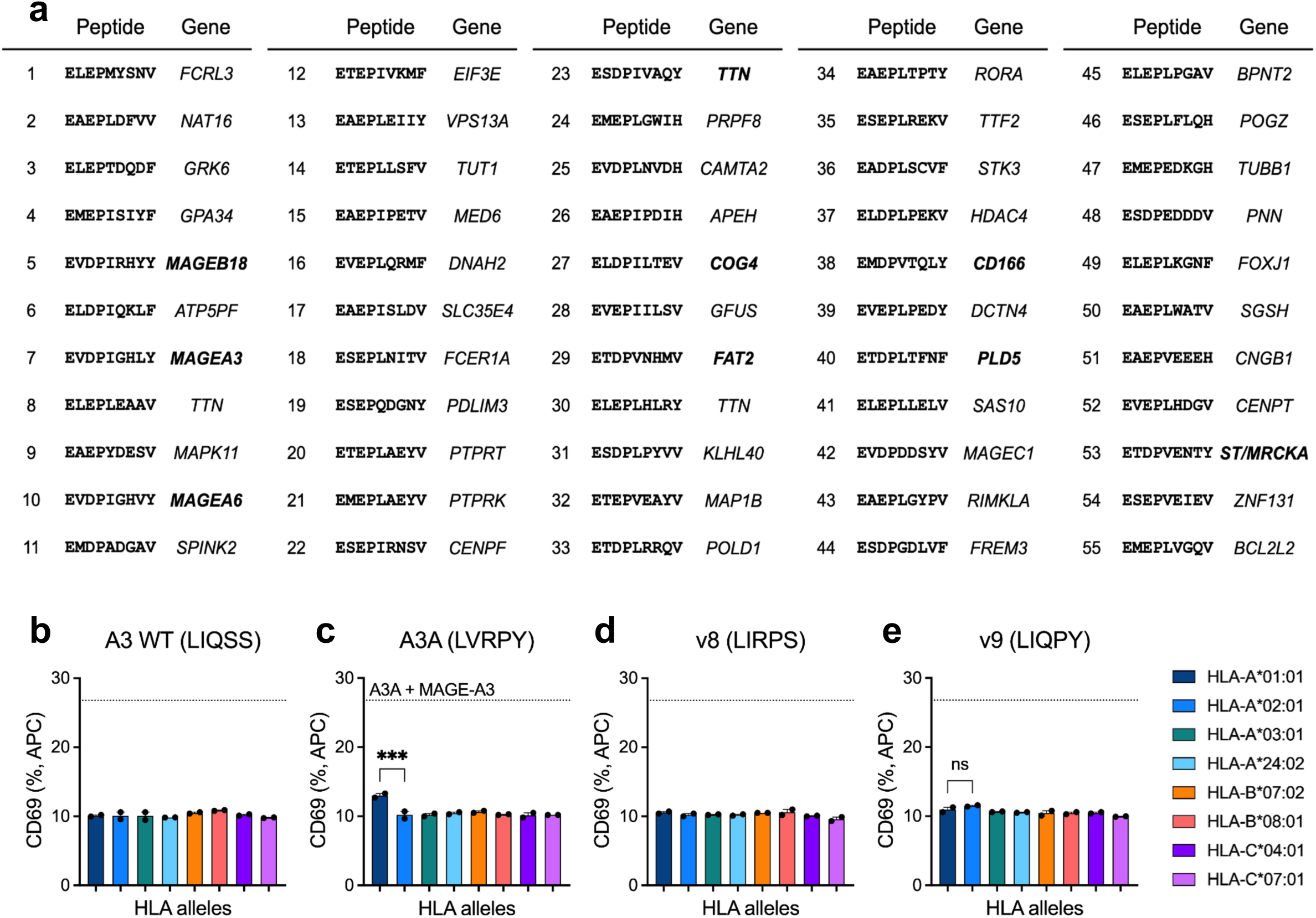
Validation of predicted v9 agonists from the 9-mer-A*01:01 yeast-display screen. **a**, The top 55 algorithm-predicted peptides from the human proteome, based on v9’s pMHC yeast-display screen, were selected for subsequent validation in co-culture assays. Peptides displayed in order of prediction ranking. Known agonists of the A3A highlighted in bold. **b-e,** K562s were transduced with eight MHC class 1 alleles (HLA-A*01:01, HLA-A*02:01, HLA-A*03:01, HLA-A*24:02, HLA-B*07:02, HLAB* 08:01, HLA-C*04:01, HLA-C*07:01) and were cultured at an E/T ratio of 1:1 for 18h with primary CD8+ T cells transduced with the TCRs **(b)** A3 WT, **(c)** A3A, **(d)** v8, **(e)** v9. Finally, anti-CD69 staining was performed on the four A3A-variant CD8+ primary T cell lines, enabling measurement of CD69 expression levels by flow cytometry. The top dotted lines correspond to the CD69 expression levels of A3A-primary CD8+ T cells after 18h of culture with A1-K562s pulsed with 100 μM MAGE-A3. Error bars represent means of technical duplicates ± SEM, and the data are representative of three independent experiments. Statistical analysis by ordinary one-way ANOVA, ***p≤0.001, ns p>0.05.

**Extended Data Fig. 10.**
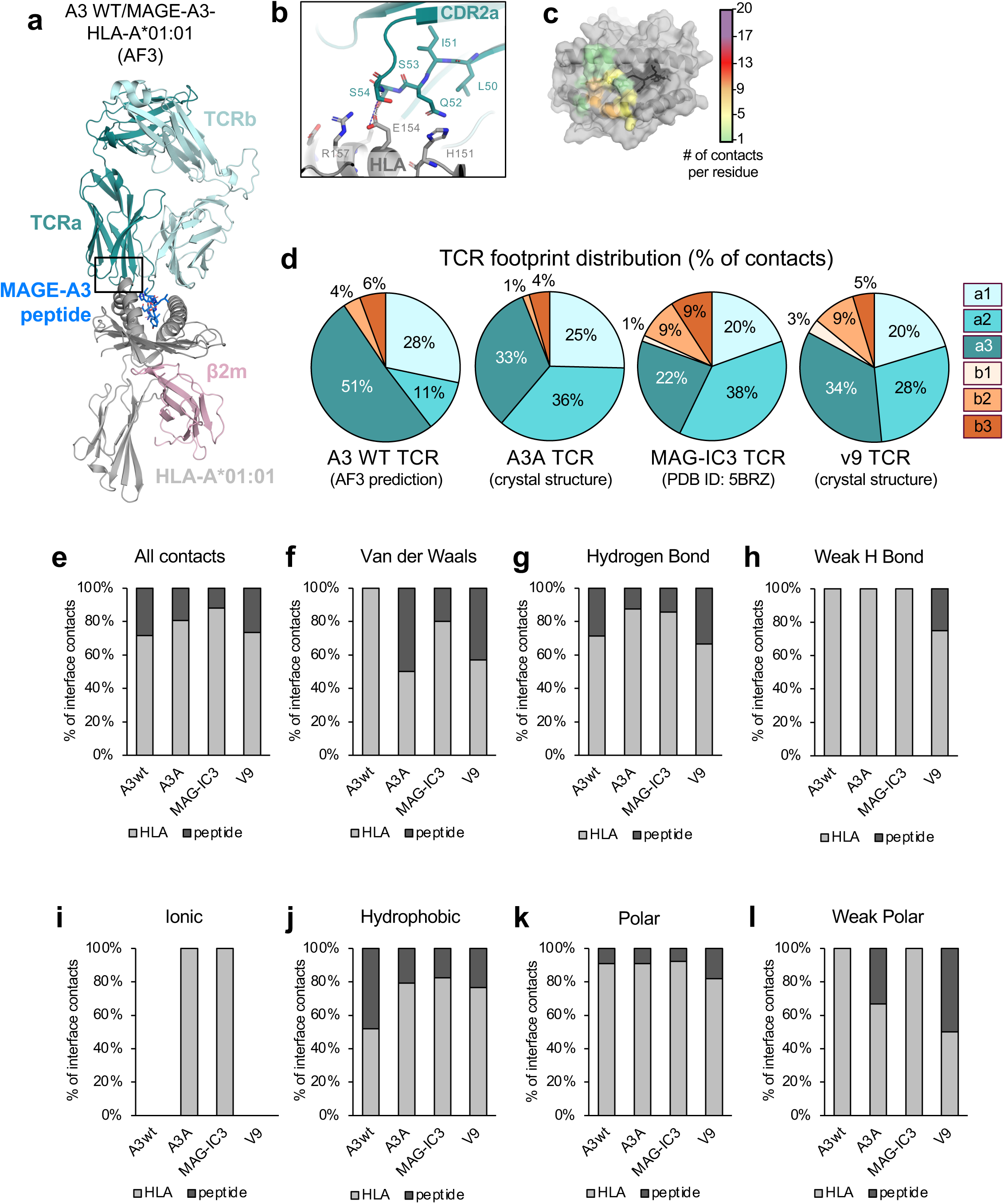
Structural prediction of the A3 WT/MAGE-A3-HLA-A*01:01 complex and comparison of the TCR/pMHC interactions by contact type. **a**, AlphaFold3 (AF3) prediction of the the A3 WT/MAGE-A3-HLA-A*01:01 complex. TCR is shown in teal, peptide in blue, β2m in pink and HLA-A*01:01 in grey. The predicted AF3 structure had high confidence values (pTM = 0.82, iPTM = 0.8) and high similarity to the A3A/MAGE-A3-HLA-A*01:01 crystal structure (r.m.s.d 0.53 Å). **b,** Close-up view of the TCR/CDR2α-HLA-A*01:01 interface, corresponding the to the black rectangle in (A). **c,** Surface representation of the MAGE-A3-HLA-A*01:01 complex, shown in grey (HLA) and dark grey (peptide). The interaction footprint of the A3 WT is shown in colour. Interacting residues on the MAGE-A3-HLA-A*01:01 surface are coloured depending on how many contacts are formed with the TCR. No contacts (grey), 1-4 (green), 5-8 (yellow), 9-12 (orange), 13-16 (red), 16-20 (purple). **d,** Contribution of CDR loops towards the TCR/MAGE-A3-HLA-A*01:01 interface, displayed as a percentage of all the contacts formed between the respective TCRs and MAGE-A3-HLA-A*01:01. **e-l,** The Arpeggio Server was used to identify all contacts that contribute to the TCR/pHLA interface, using the A3A, v9 and MAG-IC3 crystal structures, along with the AF3 prediction of the A3 WT complex. Of all the contacts, we calculated the percentage contribution of TCR/peptide (dark grey) vs TCR/HLA interactions (light grey). **(e)** represents all contacts, while **(f-l)** represent the contacts of a specific category. The interaction types are described on the Arpeggio server.

**Extended Data Fig. 11.**
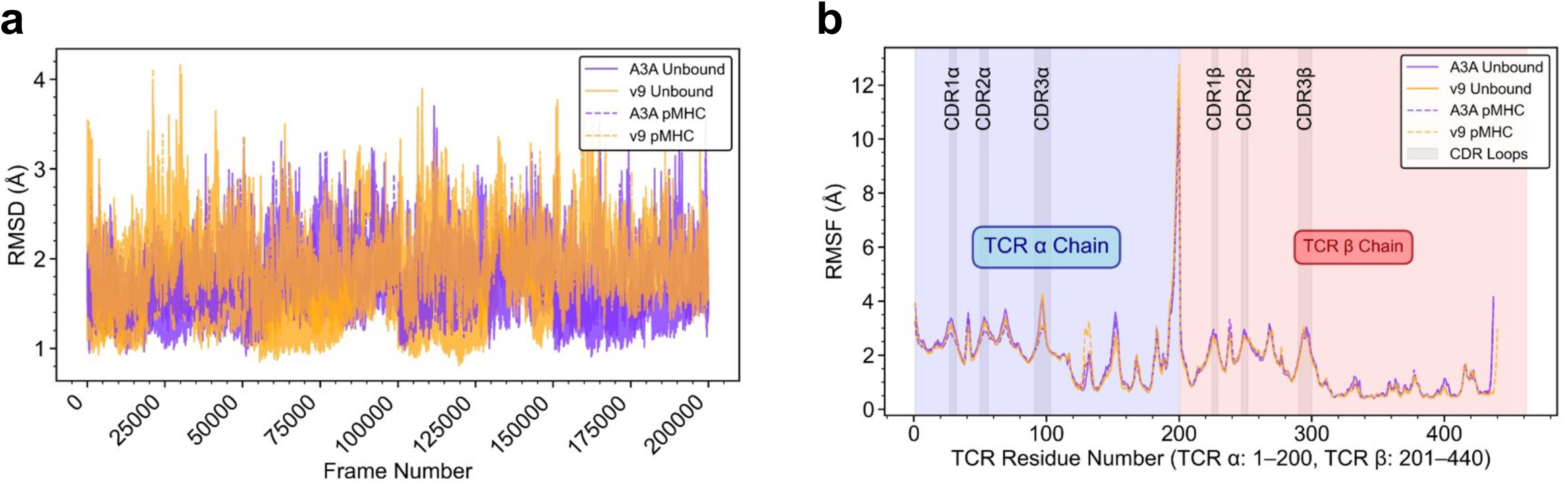
Molecular dynamics analysis of loop conformational dynamics identifies loops and interactions which show consistencies between A3A and v9. **a**, Root mean square deviation (RMSD) analysis of TCR backbone atoms over the simulation trajectory for A3A and v9 TCRs in both unbound (purple and orange solid lines, respectively) and pMHC-bound states (purple and orange dashed lines, respectively). **b,** Root mean square fluctuation (RMSF) analysis by residue number for A3A unbound (purple solid line), v9 unbound (orange solid line), A3A bound (purple dashed line), and v9 bound (orange dashed line). Shaded regions indicate CDR loops (CDR1α, CDR2α, CDR3α for residues 1-200 representing the TCR α chain; CDR1β, CDR2β, CDR3β for residues 201-440 representing the TCR β chain). RMSF values are shown in Angstroms.

**Extended Data Fig. 12.**
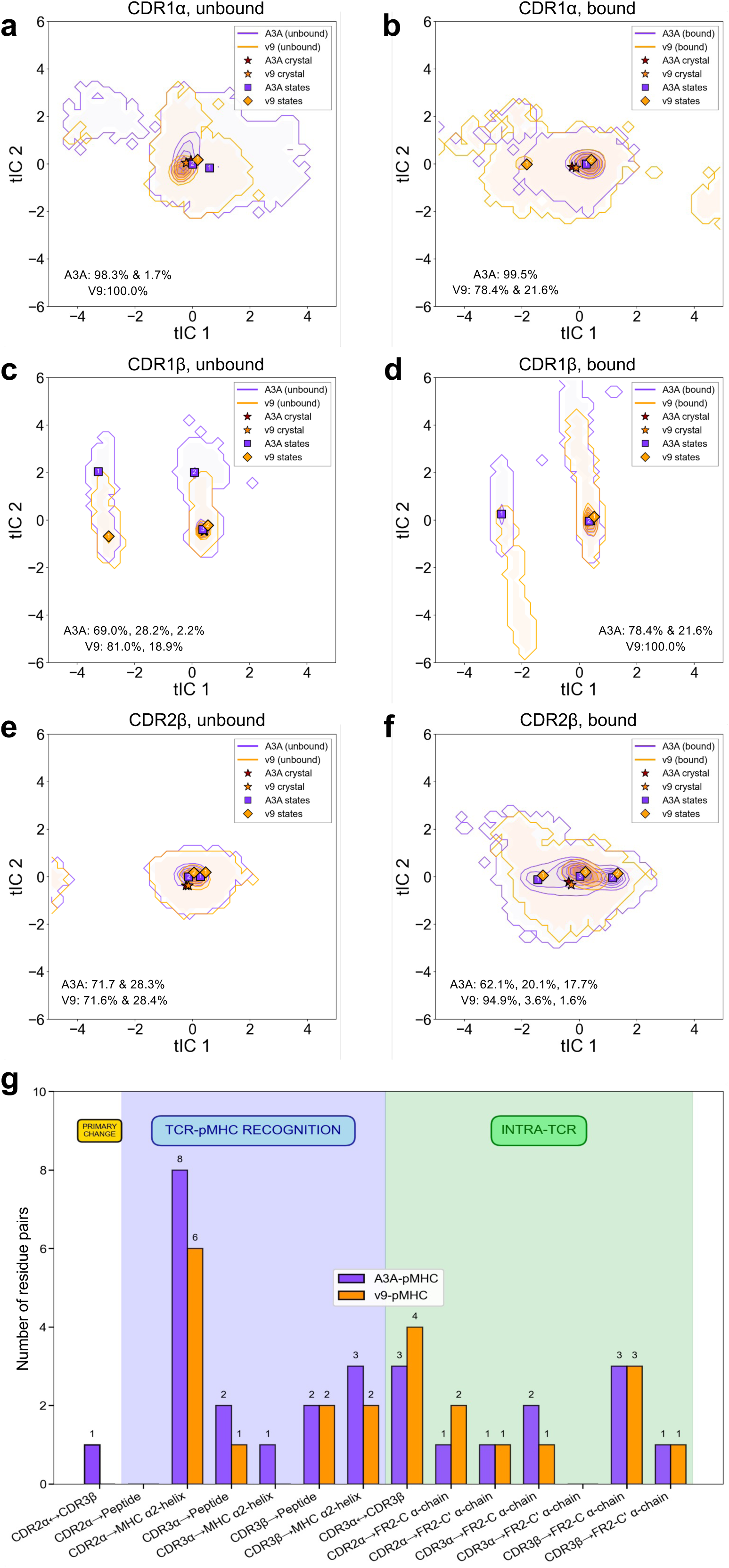
Molecular Dynamics analysis of loop conformational dynamics identifies loops and interactions which show consistencies between A3A and v9. **a-f**, Conformational overlap analysis for remaining CDR loops not shown in Figure 5. CDR1α **(a,b)** and CDR1β **(c,d)** are shown in unbound and bound states respectively, while CDR2β **(e,f)** is shown in unbound and bound states, respectively. **g,** Contact network comparison: bar chart showing number of residue pair interactions (≥25% persistence, 4.5Å cutoff) between A3A/pMHC (purple) and v9/pMHC (orange). The bars represent different contact categories: primary change (CDR2α↔CDR3β), TCR-pMHC recognition contacts (CDR loops to pMHC), and intra-TCR contacts.

**Extended Data Fig. 13.**
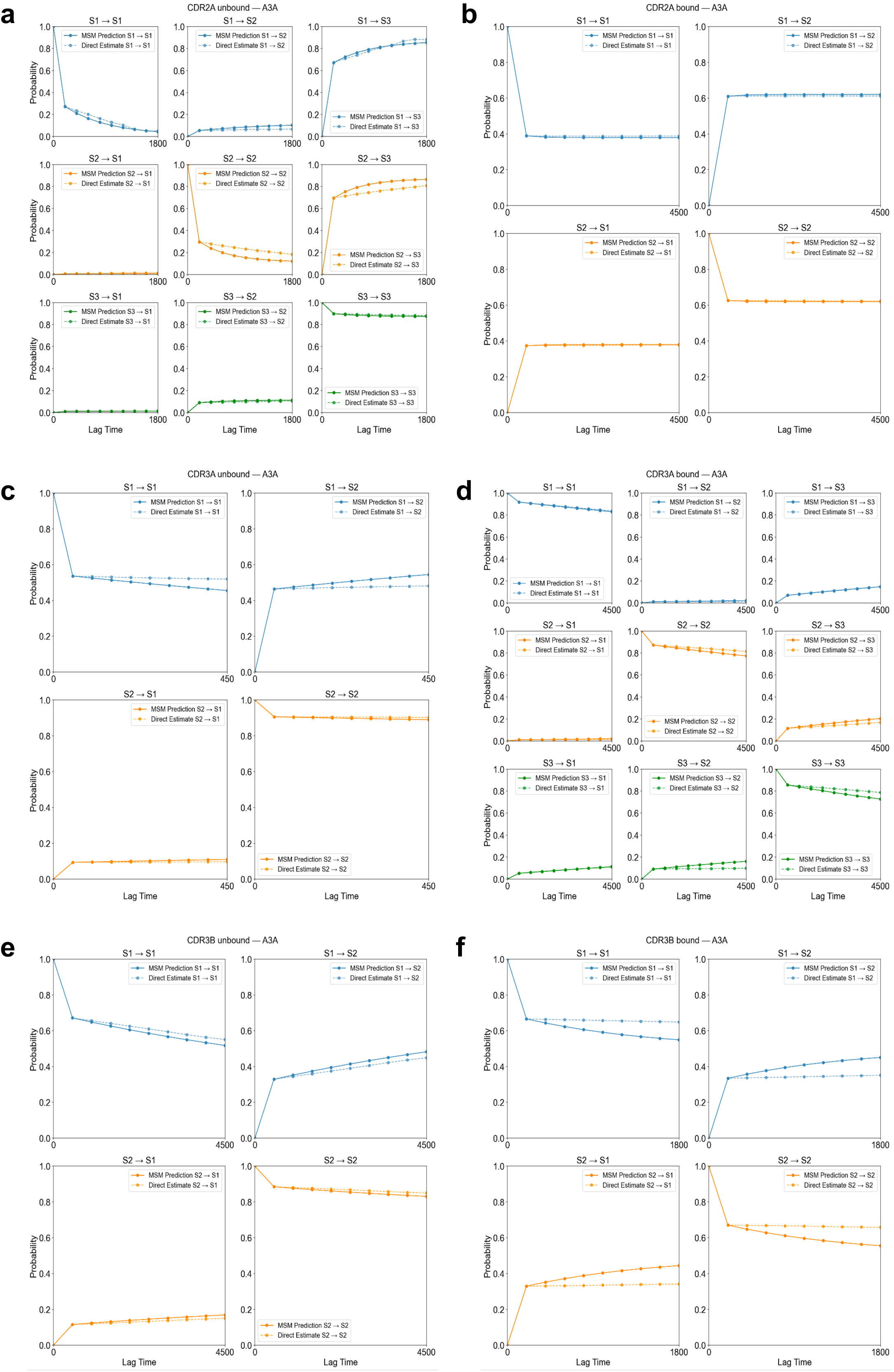
Chapman-Kolmogorov test validation for A3A MSMs. Chapman-Kolmogorov tests for: CDR2α **(a)** unbound and **(b)** bound, CDR3α **(c)** unbound and **(d)** bound, and CDR3β **(e)** unbound and **(f)** bound. MSMs for A3A. Each panel shows MSM-predicted transition probabilities (solid lines) against direct MD estimates (dashed lines) across multiple lag times for all pairwise transitions between metastable states (S1–S3). Agreement between predicted and estimated values confirms that the MSMs accurately describe the conformational dynamics at the chosen lag times.

**Extended Data Fig. 14.**
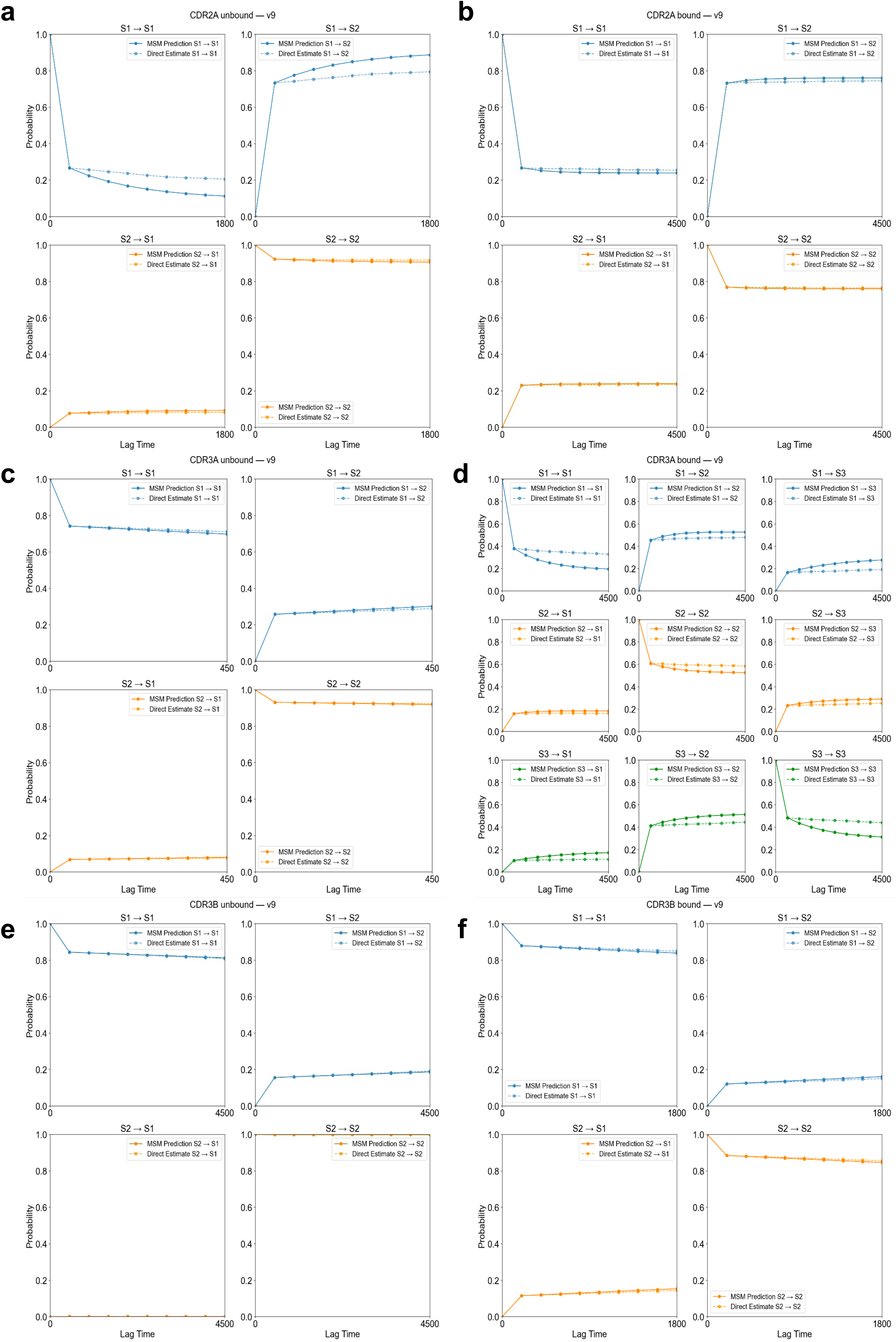
Chapman-Kolmogorov test validation for v9 MSMs. As in Extended Data Figure 13, Chapman-Kolmogorov tests for: CDR2α **(a)** unbound and **(b)** bound, CDR3α **(c)** unbound and **(d)** bound, and CDR3β **(e)** unbound and **(f)** bound. MSMs for v9.

**Extended Data Fig. 15.**
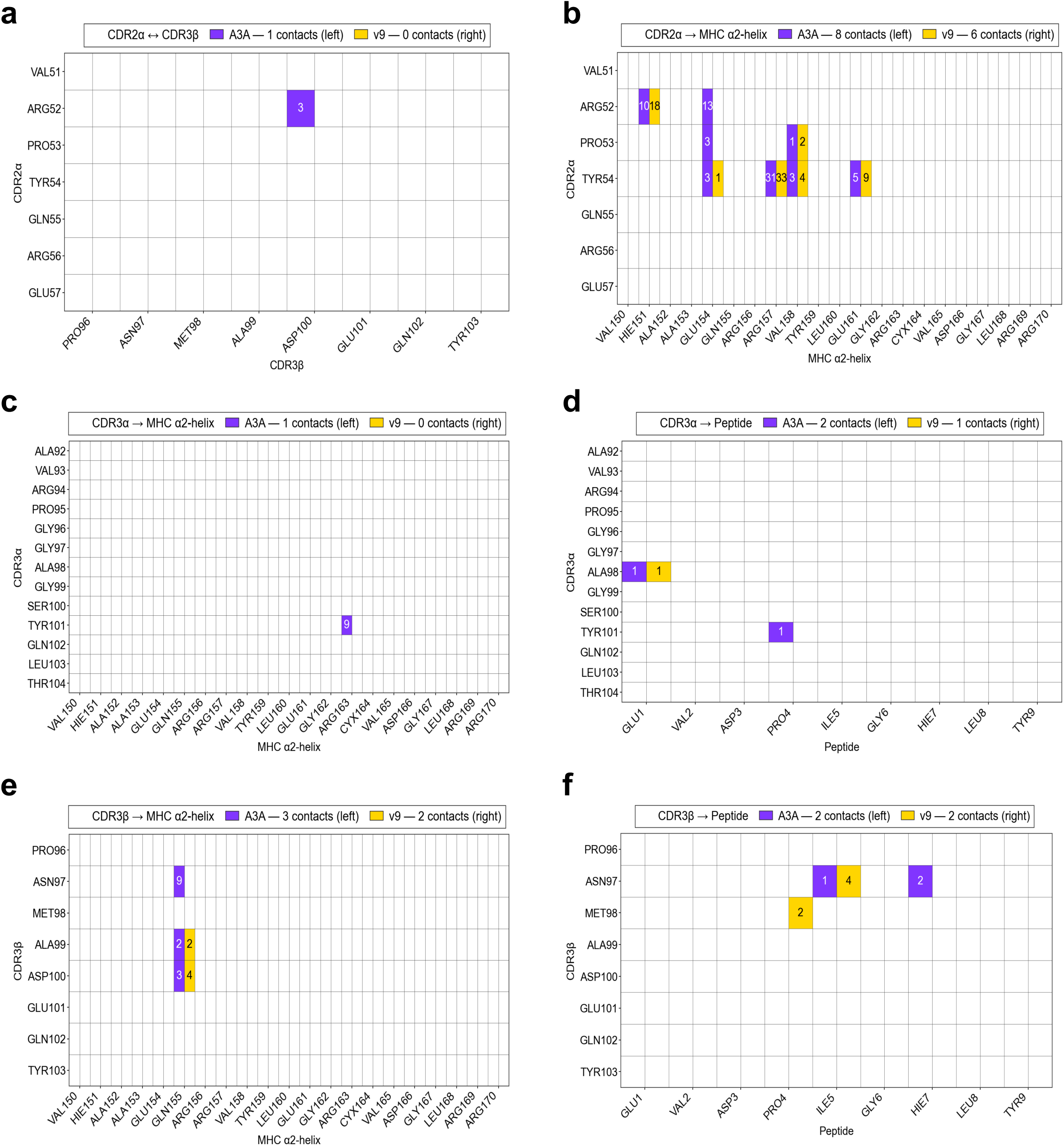
Persistent residue-pair contacts (within 4.5 Å, side-chain only, ≥25% simulation persistence) between CDR loops of A3A and v9 TCRs and the MAGE-A3 peptide or HLA-A*01:01 α2-helix. Each heatmap shows A3A contacts (purple, left half) and v9 contacts (yellow, right half), with numbers indicating atom-pair contacts per residue pair. **a,** CDR2α–CDR3β intra-TCR contacts, with the Arg52–Asp100 contact present in A3A but absent in v9. **b,** CDR2α–MHC α2- helix contacts (A3A: 8, v9: 6), with losses primarily at Arg52 and Tyr54. **c,** CDR3α–MHC α2-helix contacts, with the sole Tyr101–Arg163 contact present only in A3A. **d,** CDR3α–peptide contacts (A3A: 2, v9: 1), with Ala98–Glu1 conserved and Tyr101–Pro4 present only in A3A. **e,** CDR3β–MHC α2-helix contacts (A3A: 3, v9: 2), with Asp100–Gln155 and Ala99–Gln155 conserved and Asn97–Gln155 present only in A3A. **f,** CDR3β–peptide contacts (A3A: 2, v9: 2), with Asn97–Ile5 conserved, Asn97–His7 present only in A3A, and Met98–Pro4 present only in v9.

**Supplementary Table 1.** Crystallographic Data Collection and Refinement Statistics. Numbers in parentheses refer to the highest resolution shell. NA: Sodium ions, NAG: N-Acetyl-D-Glucosamine, EDO: Ethylene Glycol, R.m.s.d.: root mean square deviation.

|  | v9 TCR<br>+ MAGE-A3 in HLA-A*01:01 | A3A TCR<br>+ MAGE-A3 in HLA-A*01:01 |
| --- | --- | --- |
| PDB code | **29IT** | **29IU** |
| Data collection |  |  |
| Source | DLS I04 | DLS I04 |
| Wavelength $\lambda$ (Å) | 0.9537 | 0.9537 |
| No. of crystals | 1 | 1 |
| Resolution (Å) | 61.27 – 2.42 (2.46 – 2.42) | 68.95 – 2.83 (2.88 – 2.83) |
| Space group | P1 | C 1 2 1 |
| Cell dimensions |  |  |
| $a, b, c$ (Å) | 48.86, 89.55, 126.22 | 172.96, 49.17, 140.92 |
| $\alpha, \beta, \gamma$ (°) | 104.58, 91.63, 105.71 | 90, 101.87, 90 |
| Unique reflections | 74892 (3694) | 28261 (1417) |
| Multiplicity | 3.62 (3.38) | 6.86 (7.14) |
| Completeness (%) | 98.70 (97.49) | 99.95 (100.00) |
| $R_{\text{merge}}$ (%) | 0.0964 (1.2621) | 0.202 (2.354) |
| $R_{\text{meas}}$ (%) | 0.114 (1.504) | 0.219 (2.539) |
| $R_{\text{PIM}}$ (%) | 0.060 (0.808) | 0.083 (0.945) |
| $CC_{1/2}$ (%) | 0.9394 (0.4608) | 0.9952 (0.2973) |
| Average $I/\sigma$ (I) | 12.28 (0.65) | 8.10 (0.77) |
| Refinement statistics |  |  |
| Resolution (Å) | 2.42 | 2.83 |
| $R_{\text{work}} / R_{\text{free}}$ | 0.2144/0.2530 | 0.2112/0.2546 |
| No. of atoms / B factors (Å<sup>2</sup>) |  |  |
| *Protein* | 13179 / 81.79 | 6561 / 97.52 |
| *Water* | 23 / 45.11 | 12 / 54.03 |
| *NA* | 2 / 52.81 | 1 / 54.36 |
| *NAG* | 56 / 96.47 | 28 / 107.12 |
| *EDO* | 16 / 67.77 | - / - |
| R.m.s.d. bond lengths (Å) | 0.0016 | 0.0017 |
| R.m.s.d. bond angles (°) | 0.46 | 0.44 |
| Ramachandran |  |  |
| *Favored* (%) | 98.71 | 98.89 |
| *Allowed* (%) | 1.29 | 1.11 |
| *Outliers* (%) | 0.00 | 0.00 |
| Molprobability Score / percentile | 0.96 | 1.02 |

**Supplementary Table 2.** Structural features of the TCR/pMHC interfaces. The Arpeggio Server was used to identify all contacts that contribute to the TCR/pMHC interface, using the A3A, v9 and MAGIC3 crystal structures, along with the AF3 prediction of the A3 WT complex. R.m.s.d. calculations were done in PyMOL. Interface geometry values were determined using the 3DTCR server.

|  | TCR/pMHC interface structural features | A3A TCR<br>+ MAGE A3 in HLA-A*01:01 (crystal structure) | v9 TCR<br>+ MAGE A3 in HLA-A*01:01 (crystal structure) | MAG-IC3<br>+ MAGE A3 in HLA-A*01:01 (PDB ID 5BRZ) | A3 WT TCR<br>+ MAGE A3 in HLA-A*01:01 (AF3 prediction) |
| --- | --- | --- | --- | --- | --- |
| Interface geometry | r.m.s.d. (Å) | - | 0.260 (vs A3A) | 0.588 (vs A3A) | 0.531 (vs A3A) |
|  | Docking angle (°) | 56.03 | 56.85 | 57.03 | 60.20 |
|  | Incident angle (°) | 9.96 | 9.29 | 8.06 | 3.38 |
|  | TCR shift Avg X | 0.40 | 0.51 | 0.05 | 1.39 |
|  | TCR shift Avg Y | 6.94 | 6.73 | 6.42 | 4.68 |
|  | Total TCR-pMHC interface area (Å²) | 1028.4 | 1017.8 | 1026.6 | 1125.5 |
| TCR-pMHC number of contacts | Total contacts | 67 | 64 | 76 | 53 |
|  | By type |  |  |  |  |
|  | Van der Waals | 6 | 7 | 5 | 2 |
|  | Hydrogen bonds | 8 | 6 | 7 | 7 |
|  | Weak H bonds | 3 | 8 | 5 | 3 |
|  | Ionic | 4 | 0 | 4 | 0 |
|  | Hydrophobic | 29 | 30 | 34 | 25 |
|  | Polar | 11 | 11 | 13 | 11 |
|  | Weak Polar | 6 | 2 | 8 | 5 |
|  | By peptide vs MHC |  |  |  |  |
|  | Peptide contacts | 13 | 17 | 9 | 15 |
|  | MHC contacts | 54 | 47 | 67 | 38 |
|  | By TCR CDR chain |  |  |  |  |
|  | a chain CDR1/2/3 | 17/24/22 | 13/18/22 | 15/29/17 | 15/6/27 |
|  | b chain CDR1/2/3 | 0/1/3 | 2/6/3 | 1/7/7 | 0/2/3 |

**Supplementary Table 3.** Full residue-pair contact analysis for A3A and v9 TCRs across all CDR loops, quantified in both crystal structures and MD simulations using a 4.5 Å sidechain heavy-atom distance cutoff. MD contacts represent residue pairs with ≥25% persistence across the trajectory. Contact types include interactions between CDR loops (CDR1α/β, CDR2α/β, CDR3α/β) and the peptide, MHC α1-helix or MHC α2-helix, as well as intra-TCR CDR–CDR contacts. Purple columns indicate A3A data and orange columns indicate v9 data. The reduction in contacts between crystal structure and MD simulation reflects filtering of transient interactions, while the consistent reduction from A3A to v9 across both methods confirms a genuinely sparser binding interface in v9.

| Contact Type | A3A Crystal | V9 Crystal | A3A MD | V9 MD |
| --- | --- | --- | --- | --- |
| CDR2α ↔ CDR3β | 1 | 0 | 1 | 0 |
| CDR3α ↔ CDR3β | 3 | 4 | 3 | 4 |
| CDR1α → Peptide | 3 | 3 | 2 | 2 |
| CDR1α → MHC α1-helix | 0 | 0 | 0 | 0 |
| CDR1α → MHC α2-helix | 4 | 4 | 2 | 1 |
| CDR2α → Peptide | 0 | 0 | 0 | 0 |
| CDR2α → MHC α1-helix | 0 | 0 | 0 | 0 |
| CDR2α → MHC α2-helix | 8 | 8 | 8 | 6 |
| CDR3α → Peptide | 2 | 2 | 2 | 1 |
| CDR3α → MHC α1-helix | 2 | 2 | 1 | 1 |
| CDR3α → MHC α2-helix | 3 | 3 | 1 | 0 |
| CDR1β → Peptide | 1 | 1 | 1 | 0 |
| CDR1β → MHC α1-helix | 0 | 0 | 0 | 0 |
| CDR1β → MHC α2-helix | 0 | 0 | 0 | 0 |
| CDR2β → Peptide | 1 | 1 | 0 | 0 |
| CDR2β → MHC α1-helix | 3 | 3 | 0 | 0 |
| CDR2β → MHC α2-helix | 0 | 0 | 0 | 0 |
| CDR3β → Peptide | 4 | 3 | 2 | 2 |
| CDR3β → MHC α1-helix | 0 | 0 | 0 | 0 |
| CDR3β → MHC α2-helix | 4 | 2 | 3 | 2 |
| TOTAL | 39 | 36 | 26 | 19 |

**Supplementary Table 4.** A3A wild-type agonist peptide predictions. The predicted A3A wild-type peptides following application of an *in silico* motif similarity analysis *(27)* to the A3A yeast display deep sequencing data (Fig. 1c). The top 55 predictions were validated in Extended Data Fig 6.

**Supplementary Table 5.** Refinement of A3A wild-type agonist peptide predictions. Refined predictions of the A3A wild-type peptides following application of an agonist-trained *in silico* motif similarity analysis *(27)* to the A3A yeast display deep sequencing data (Fig. 1c). The top 55 predictions were validated in Extended Data Fig. 6.

**Supplementary Table 6.** v9 wild-type agonist peptide predictions. The predicted v9 wild-type peptides following application of an *in silico* motif similarity analysis *(27)* to the v9 yeast display deep sequencing data (Fig. 1C). The top 55 predictions were validated in Extended Data Fig. 9.

## Materials and methods

### Production of a peptide-HLA-A*01:01 scaffold for yeast display

For the MAGE-A3-A*01:01 construct, the MAGE-A3 peptide DNA sequence was annealed by PCR upstream of a β2M-A*01:01-c-Myc DNA fragment synthesised by IDT. All cloned DNA plasmids in this manuscript were validated by Sanger sequencing to confirm sequence integrity prior to downstream application.

Yeast scaffolds were generated by electroporation of competent EBY100 cells (treated with Tris-DTT and frozen in E buffer) with the cloned MAGE-A3-A*01:01-pYAL plasmid, as described previously^9^. In short, 1 μL of plasmid at 200-500 ng/μL was added to approx. 5 × 10^7^ competent EBY100 cells for 5 minutes on ice. The cells were transferred to electroporation cuvettes and electroporated using a MicroPulser (Bio-Rad 1652100) at 1.50 kV with an expected time constant of 3.00-5.00 ms. Electroporated EBY100 cells were recovered using 3 mL of the selective glucose (dextrose) growth medium (SDCAA) and incubated at 270 rpm at 30°C for 48h or until confluent (OD>5). Once confluent, the yeast were passaged into SDCAA overnight, then transferred to a selective galactose growth medium (SGCAA) to achieve an OD of 0.5, and incubated at 270 rpm at 20°C for 48h to allow expression of the MAGE-A3-A*01:01 gene and display on the yeast cell surface. Flow cytometric validation of the yeast displayed peptide-HLA-A*01:01 complex was conducted as follows: 5 × 10^6^ yeast cells were stained in 75 μL MACS buffer (PBS, BSA, EDTA, 0.09% NaN3) with an AF647 anti-myc tag antibody at a 1/150 concentration (Cell Signalling, cat no. 2233S), and a TCR tetramer with 1 μM PE Streptavidin (BioLegend, cat no. 405204) prepared at a 5:1 TCR:SA ratio, using an Attune CytKick Autosampler (ThermoFisher) and analysed using FlowJo software version 10.9.0.

### Generation of a combinatorial 9-mer-HLA-A*01:01 library for yeast display

To generate a combinatorial 9-mer library with high diversity, all peptide positions (excluding the anchors) were randomised. To optimise peptide binding to HLA-A*01:01 scaffold, peptide anchor positions at P3 and P9 were limited to Glu-Asp and Tyr. To achieve a balanced codon distribution while avoiding stops, the trimer phosphoroamidites technology was utilised for oligo synthesis. Specifically, oligos with a subset of trimer phosphoroamidites were ordered to avoid stops and cysteines (iTriMix19), amounting to 19 possible codons at each peptide position. The theoretical 9-mer diversity was therefore 19^7^ × 2(p3) × 1(p9), summing to 1.8 × 10^9^ possible 9-mers. Codon-balanced iTriMix19 9-mer oligos were synthesised by IDT. The random 9-mer sequences and GlySer linker were inserted upstream of the β2M-A*01:01-c-Myc construct by PCR, and the fully formed construct was then inserted into the digested pYal-Aga2p vector. The amplification process was conducted on a Bio-Rad T100 Thermal Cycler using a standard ITA buffer composition and reaction scheme. Agarose gels were subsequently used to extract the amplified PCR products at the correct size and purity.

Electroporation of the randomised 9mer-A*01:01 library was conducted as follows. Two 5 mL EBY100 cultures were inoculated in YPD and grown overnight. The following day, both yeast cultures were passaged in sufficient YPD for 8L of confluent culture the next day. The following morning, the OD600 of the overnight cultures was measured, enabling a new starting culture of 300 mL YPD at OD600=0.3 in a 1 L baffled flask that was left to grow until OD600=1.6. Next, 3 mL of Tris/DTT (1 M Tris pH 8.0, 2.5 M 1,4-dithiothreitol, filter-sterilised) plus 15 mL of 2M LiAc/TE were added to the yeast culture and incubated shaking for 15 min. The yeast were then pelleted at 3500g for 3 min, 4°C, and resuspended in 50 mL ice-cold NewE buffer (10 mM Tris pH 7.5, 270 mM sucrose, 1 mM MgCl2, filter-sterilised). This was repeated, before a third resuspension now in 10 mL ice-cold NewE buffer. After the final spin, the pellet was resuspended in 600 μL E buffer + 50 μg 9mer-HLA-A*01:01 insert + 10 μg digested pYal plasmid, and 150 μL of the yeast/DNA mix was distributed into 2mm gap electroporation cuvettes for electroporation at 2500V using a Bio-Rad electroporator (time constants between 2.9-4 ms1). The yeast were recovered in 200 mL YPD for 1 h, prior to transfer into 330 mL SDCAA for selective growth overnight. If the yeast reached an OD600=5 the next day, the culture was passaged into 300 mL fresh SDCAA at OD=0.3 overnight, and induced the following morning in log phase in 330 mL SGCAA + 5% SDCAA for 48h at 20°C, ready for TCR selections. Flow cytometric validation was conducted as described above.

### Soluble TCR expression in Expi293 mammalian expression system

The TCR α- and β- variable DNA sequences were synthesised by IDT and cloned into appropriate backbones by Gibson Assembly, with the respective TCR constant region DNA sequences previously amplified by PCR. The destination vectors were digested with appropriate restriction enzymes: pD649 Not1 (NEB R3189) and BamH1 (NEB R3136). TCR α-chains were cloned into the pD649 expression vector encoding a C-terminal acidic helix zipper and a BirA biotinylation tag, and β-chains were cloned into the same vector encoding a C-terminal basic helix zipper.

Transfection and expression in the Expi293 system was conducted as per the manufacturer’s instructions (Expi293 Expression System, ThermoFisher Scientific). In short, 50 mL of Expi293F cells were co-transfected with 25 µg of the variant TCRα pD649 plasmid and 25 µg of the a3a TCRβ pD649 plasmid (gene fragments ordered from Twist), alongside 150 µL of Expifectamine reagent for a final DNA quantity of 50 µg at 1 µg/mL, at a density of 3 × 10^6^ cells/mL. At 18h post-transfection, transfection enhancers 1 and 2 were added to the cells as per the manufacturer’s protocol. On day 5 post-transfection, the supernatant containing soluble TCR protein was harvested and the Expi293F cells were removed by centrifugation at 3,200g for 10 minutes at 4°C. The supernatant was then diluted two-fold in 1x HBS buffer (30 mM HEPES pH 7.4, 150 mM NaCl), conditioned by adding 20 mM Tris-HCl pH 8.0 and 0.05% (v/v) NaN_3_, and incubated with 3 mL of 50% Ni-NTA agarose bead suspension (Qiagen 30210) overnight at 4°C with gentle stirring throughout.

Recombinant polyHis-tagged TCR protein was purified using IMAC. The IMAC eluate was then concentrated to a final volume of ∼500 µL. The purified TCRs were subsequently biotinylated in preparation for downstream application as a MAGE-A3-A*01:01 yeast display detection probe; purified TCRs were incubated with recombinant *E.coli* BirA (SinoBiological, 5 µg BirA per 10 nmol of protein) in reaction buffer consisting of 50 mM bicine pH 8.3, 10 mM ATP, 10 mM magnesium acetate and 50 µM D-Biotin, at room temperature for 60 minutes and then overnight at 4°C.

The next day, biotinylated TCR proteins were purified via size-exclusion chromatography (SEC) on an AKTA Pure system fitted with a Superdex 200 10/300 GL Increase column equilibrated with 1x HBS, at a flow rate of 0.7 mL min^-1^. The fractions corresponding to the appropriate peaks on the elution profile (12 mL retention volume for biotinylated αβ TCRs) were collected and analysed using SDS-PAGE. Fractions that resulted in bands of appropriate size and purity were then pooled and concentrated using Amicon Ultra-4 Centrifugal Filter Units (Sigma-Aldrich UFC8100, ≤100 kDa MWCO) to 1-3 mg/mL. The purified, biotinylated TCRs were then aliquoted and flash-frozen in liquid nitrogen for storage at -80°C.

### Selection of the 9-mer-HLA-A*01:01 library with recombinant TCRs

All selections were conducted as described previously ^9^ using biotinylated soluble TCR coupled to streptavidin-coated magnetic MACS beads (Streptavidin MicroBeads, Sab; Miltenyi; cat no. 130-048-101). In short, 10× diversity of 9-mer-A*01:01 yeast were negatively selected with 250 µL SAb for 1 hr rotating at 4°C in 10 mL of PBS + 0.5% bovine serum albumin and 1 mM EDTA (PBE). Yeast were pelleted and passed through an LS column (Miltenyi; cat no. 130-042-401) attached to a magnetic stand (Miltenyi; cat no. 130-042-303) and washed with 10 mL PBE three times. The flowthrough was then incubated for 2hr rotating at 4°C with 250 µL SAb pre-incubated with 400 nM biotinylated TCR for 15 minutes in the dark at 4°C. Once again, yeast were pelleted and passed through an LS column, and the elution was grown in 3 mL SDCAA pH 4.5 overnight in 14 mL vented culture tubes after one 5 mL SDCAA wash. Once yeast reached OD600>5, they were induced in SGCAA pH 4.5 with 5% SDCAA for 48h prior to the next selection round. All subsequent selections (rounds 2-4) were conducted with 50 µL SAb or TCR-coated SAb in 450 µL of PBE, and washed with 3 mL PBE. All yeast grown in 14 mL vented culture tubes were shaken at 270 rpm.

Each round was monitored post-induction with anti-c-Myc staining. Further, once all selection rounds were complete, the naïve library and all subsequent rounds were stained with an anti-c-Myc antibody AF647 (Cell Signalling; cat no. 2233S) and 1 µM SA-PE (BioLegend; cat no. 405204) TCR tetramer (5:1 TCR:SA) for 1h at 4°C in MACS buffer.

### Extraction of the TCR-selected 9-mer peptides for deep sequencing analysis

9-mer-A*01:01 constructs were isolated from 5 × 10^7^ yeast per round of selection by miniprep (Zymoprep II kit Zymo Research). Deep sequencing was conducted using an Illumina Novaseq X Plus platform. To prepare the samples, two 25 cycle PCR amplifications were conducted, generating sequences containing: Illumina P5-TruSeq read 1-(N8)-Barcode-pHLA-(N8)-TruSeq read 2-Illumina P7. All PCR products following both amplification steps were purified by agarose gel purification and quantified by nanodrop. The samples were subsequently deep sequenced by Novogene.

**Step 1**: the Illumina chip primer sequences and random 4bp sequences were added to the flanking regions of the sequencing product using “Illumina Nextera Forward primer” and “Illumina Nextera Reverse primer”.

**Illumina Nextera Forward primer:** 5’TCGTCGGCAGCGTCAGATGTGTATAAGAGACAGNNNNGCTGTTTTTCAATATTTTCTGTTATTGCTAGCG 3’

**Illumina Nextera Reverse primer:** 5’GTCTCGTGGGCTCGGAGATGTGTATAAGAGACAGNNNNAACTTGGATCTTTGGAGTTCTTTGTATGGA 3’

**Step 2**: barcodes/indices were added upstream and downstream of the sequencing product using unique “i5 Forward primers” (unique indices in Table 2) and “i7 Reverse primers” (unique indices in Table 3).

**i5 Forward primer** (*example barcode in italics*, 8bp barcode indices in Table 2)

5’ AATGATACGGCGACCACCGAGATCTACAC*TAGATCGC*TCGTCGGCAGCGTC 3’

**Table 2.** Forward primer i5 indices, 5’.

|  |  |  |  |
| --- | --- | --- | --- |
| N501 | TAGATCGC | N505 | GTAAGGAG |
| N502 | CTCTCTAT | N506 | ACTGCATA |
| N503 | TATCCTCT | N507 | AAGGAGTA |
| N504 | AGAGTAGA | N508 | CTAAGCCT |

**Table 3.** Reverse primer i7 indices, 5’.

|  |  |  |  |
| --- | --- | --- | --- |
| N701 | TCGCCTTA | N707 | GTAGAGAG |
| N702 | CTAGTACG | N708 | CCTCTCTG |
| N703 | TTCTGCCT | N709 | AGCGTAGC |
| N704 | GCTCAGGA | N710 | CAGCCTCG |
| N705 | AGGAGTCC | N711 | TGCCTCTT |
| N706 | CATGCCTA | N712 | TCCTCTAC |

**i7 Reverse primer** (*example barcode in italics*, 8bp barcode indices in Table 3)

5’ CAAGCAGAAGACGGCATACGAGAT*TCGCCTTA*GTCTCGTGGGCTCGG 3’

### Prediction of wild-type TCR antigens using computational algorithms

Deep sequencing results were analysed following the protocol of ref. 9^9,27,31^. Paired-end reads were demultiplexed based on the unique barcodes and translated to amino acids. Sequences containing frameshifts or stop codons were removed from downstream analysis. For wild-type antigen predictions, a positional frequency matrix-based algorithm was applied to the extracted peptide data^27^.

### Positional Frequency Matrices

To infer the overall similarity across the peptides bound by a given TCR, the top 95% of peptides enriched from round 4 were retrieved and further analysed. For each position of each individual peptide, the PMBEC similarity score^26^ was calculated in relation to all other existing peptides within the 95% filter. The averages of these values were taken, which gave a score on how similar each position of a given peptide was in comparison to the rest. To retrieve a score indicating the overall similarity per position, these values were normalised by the square root of the enrichment count and averaged to retrieve one similarity score for each position. For plotting the data, a similarity matrix was generated based on the individual peptide scores and ordered using hierarchical clustering, which did not take the anchor positions into account. Finally, the sum/average of all positional similarity scores (excluding anchor residues) were calculated to give a single similarity metric for each TCR.

### Generation of secondary T cell and antigen-presenting cell lines

The pHR vector with MluI and Not1 restriction sites was used for the generation of TCR-Jurkat T and A1-K562 cell lines. For CD8 expression on Jurkat T cells, the pHRi vector was used. The TCR Vα and Vβ fragments were synthesised with the respective wild-type A3A signal peptides (IDT) with suitable bp overhangs for Gibson Assembly into separate pHR vectors alongside the full length Cα and Cβ fragments, respectively, with V and C fragments in equimolar quantities. The HLA-A*01:01 heavy chain sequence (along with all other HLA alleles) were obtained from IPD-IMGT/HLA and synthesised along with its endogenous signal peptide (IDT) for cloning into the pHR vector. The products of the TCRα, TCRβ, and HLA-A*01:01 ITA reactions were used to transform DH5α *E. Coli* (NEB C2987H) that were subsequently grown on LB agar plates with 1µg/mL carbenicillin overnight. Single bacterial colonies were grown in liquid LB bacterial culture medium with 1µg/mL carbenicillin overnight, and the cloned DNA plasmids were extracted and purified the following morning. All cloned DNA plasmids were sent for Sanger sequencing to confirm sequence integrity prior to downstream application.

Lentivirus infections were used to generate stably transfected cell lines expressing the desired genes (TCRα, TCRβ, and HLA-A*01:01 heavy chain): HEK-293T cells, grown in DMEM supplemented with 10% FBS, 1% GlutaMax, and 1% penicillin/streptomycin (v/v), were plated in six well plates with 3 × 10^5^ cells/plate (2 mL total) (all reagents from Gibco, Thermo Scientific, UK). After 24h, for each well of cells, 750 ng plasmid of interest, 500 ng psPAX, and 260 ng pMD2.G were mixed with 4.5 μL Fugene transfection reagent (Promega) in 100 μL Opti-MEM and rested for 20 minutes. In the meantime, cDMEM was removed from each well, and fresh cRPMI was added to each well. After 20 minutes, the DNA/Fugene mixture was added to each well, the plates were gently rocked up-down, side-side, and then incubated for 3 days. All cell lines grown at 37°C, 5% CO_2._ 72 hours later, the lentivirus supernatants were removed and spun down at 3300 rpm for 10 minutes to remove any unwanted cells. For CD8+ TCR Jurkat T cells, 500 μL of TCRα lentivirus and 500 μL of TCRβ lentivirus were added to 1 × 10^6^ TCR Jurkat T cells previously co-transduced with CD8α-pHRi and CD8β-pHRi vectors. For APC K562 cells, 500 μL of the HLA-A*01:01 (or other) heavy chain was added to 0.5 × 10^6^ WT K562 cells, assembling with endogenous β2M in the ER. Transduced Jurkat T and K562 cells were grown in cRPMI. All cell lines grown at 37°C, 5% CO_2._ Cells were analysed using FACS after 5 days of lentiviral infection to confirm expression of the desired proteins (anti-TCR PE, BioLegend; anti-CD8 PB, BioLegend).

### Co-culture validation of antigen predictions

Peptides (GenScript, purity >70%) were dissolved in DMSO, aliquoted, and stored at -80°C (maximum x5 freeze-thaw cycles). A1-K562 cells (split the day before) were resuspended at 1×10^6^ cells/mL, and 100 μL were aliquoted to each well of a 96-well U-bottom plate, supplemented with 100 μL fresh cRPMI. A1-K562 cells were pulsed with single concentrations of thawed peptides (<1% DMSO), or if conducting a peptide titration, the starting peptide concentration was prepared at the desired volume in a separate 96-well U-bottom plate, and titrated in thirds before transfer to the cells. The A1-K562 cells were peptide pulsed for 2 hours at 37°C, and negative controls (A1-K562 cells not pulsed with peptide) were prepared in the same way without peptide. Pulsed A1-K562 cells were then washed once to remove excess peptides (1300rpm, 3mins). TCR Jurkat T cells (split the day before) were resuspended at 5×10^5^ cells/mL and 200 μL added directly to each well of washed peptide-pulsed A1-K562 cells (i.e. a 1:1 E/T ratio). The stimulation was performed at 37°C for 14-16 hours.

Following stimulation, flow cytometric analysis was conducted to quantify cell surface protein expression. Samples of 5-10x10^5^ cells were harvested and centrifuged at 1300 rpm for 3 minutes in appropriate polypropylene U-bottom 96 well plates. Residual medium was washed off by resuspending the pellets twice in 200 μL ice-cold MACS buffer (BSA, EDTA, 0.09% azide in PBS) and centrifuging the cells again at 1300 rpm for 3 minutes. To specifically label cell surface proteins, fluorescently conjugated antibodies were used at 10 μg/mL unless otherwise indicated by the manufacturer’s protocol. Mammalian cells were stained with 50 μL of antibody mix for 45 minutes on ice in the dark (anti-CD69-APC, and anti-αβTCR-PE or anti-CD45-PE) on ice for 45 minutes, washed twice, and immediately analysed by an Attune NxT Flow Cytometer (Thermo Fisher). Excess antibody was subsequently washed off as previously described and resuspended in 130 μL of MACS buffer prior to analysis by an Attune CytKick (Thermo) using appropriate lasers. Viable cells were gated based on forward and side scatter, and fluorescence emission was collected by gates set by negative control samples. For functional assays, an additional viability dye was used (Zombie Violet) prior to cell surface antibody staining, following the manufacturer’s instructions. Raw FCS files were exported from the Attune and analysed using FlowJo software version 10.9.0.

### Generation of primary CD8+ T cell lines

For maximal CD8+ T cell transduction efficiency, TCRβ-P2A-TCRα lentiviral vector constructs were cloned and used to transfect HEK293T cells for lentivirus expression, as previously described. Large lentivirus batches were made on separate occasions, harvested, and frozen at -20°C for <1 month. On the day of transduction, lentivirus was thawed at 37°C and added directly to the cells. All cell culture of human T cells was done using cRPMI (10% FBS, 1% glutamax, 1% sodium pyruvate, 1% HEPES, 1% penicillin/streptomycin) at 37°C, 5% CO_2_.

**Day 0**: T cells were isolated from whole blood from healthy donor leukocyte cones purchased from the NHS Blood and Transplantation service at the John Radcliffe Hospital, Oxford. This project was approved by the Medical Sciences Interdivisional Research Ethics Committee, with ethics approval reference: R93485/RE001. CD8+ T cells were isolated from fresh leukocyte cones using the RosetteSep Human CD8+ enrichment cocktail (STEMCELL) at 150 μL/ml. After a 20 min incubation at room temperature, blood cone samples were diluted 3-fold with PBS and layered onto Ficoll (STEMCELL Technologies) at a 0.8:1.0 Ficoll:sample ratio, and spun at 1200g for 20 min at room temperature, acceleration 2, deceleration 0. Buffy coats were harvested with Pasteur pipettes, washed twice in cRPMI, counted, and the cells were rested for 1-4h at 1.0 × 10^6^/mL in 24 well plates (1 mL per well) with 30 U/mL IL-2 (BioLegend). The cells were then treated with CD3/CD28 Human T-activator dynabeads (Thermo Fisher) at a 1:1 bead:cell ratio for 48h for proliferation and expansion (according to the manufacturer’s instructions).

**Day 2**: after 48h, all cells were harvested, counted and resuspended at 0.5 x 10^6^/mL in cRMPI. 2 mL cell suspension was then added to each well of a 12 well plate (i.e. 1 x 10^6^ cells per well) and treated with 1.5 mL TCR lenti-virus (transduction was validated by transducing -/- Jurkat T cells with the same lenti-virus batches and confirming >95% expression levels). Each well was supplemented with 400 μL cRMPI with IL-2 at a concentration of 300 U/mL, for a final concentration of 30 U/mL (in 4 mL total volume). Two wells were also reserved for un-transduced controls.

**Day 4**: after 48h, the cells were expanded at a 1:1 ratio with cRPMI and fresh IL-2 to a final concentration of 30 U/mL.

**Day 6**: after 48h, the dynabeads were magnetically removed, and the CD8+ T cells were suspended at 1.0 x 10^6^/mL in cRPMI with fresh IL-2 to a final concentration of 30 U/mL. The cells were left overnight and used for Incucyte cytotoxicity experiments the following morning. TCR expression was confirmed by flow cytometry staining with a MAGE-A3-A*01:01-PE-conjugated tetramer at 0.2 μM SA-PE.

### Incucyte cytotoxicity assays

A375 cells were grown in adherent T75 flasks in cDMEM, DMEM supplemented with 10% FBS, 1% GlutaMax, 1% sodium pyruvate, and 1% penicillin/streptomycin (v/v), and passaged every 3 days. A1-HEK293T cells were transduced with HLA-A*01:01 lentivirus by culturing 2.0 × 10^6^ HEK293T cells (at 1.0 × 10^6^/mL) with 2 mL HLA-A*01:01 lentivirus and 6 mL fresh cDMEM in a T25 flask for 48h. Once confluent, the cells were expanded into a T75 flask and passaged every 3 days with fresh cDMEM. On the afternoon of the day before the Incucyte experiment the target cells were seeded in fresh cDMEM into flat bottom 96 well plates (in 100 μL/well) and rested overnight: (5000 A375 cells per well, or 3000 A1-HEK293T cells per well).

To prepare the cytotoxicity assays, 50 μL of media was carefully removed per well, leaving 50 μL of target cells per well. To 5000 A375 cells, 50 μL of primary CD8+ T cell lines were subsequently added at the desired concentration to achieve a 1:10 (1.0 × 10^4^/mL), 1:5 (2.0 × 10^4^/mL) and/or 1:2 (5.0 × 10^4^/mL) E/T ratio. 3000 A1-HEK293T cells were first pulsed with peptide at 50 μM for 1h prior to directly adding 50 μL of the primary CD8+ T cell lines at the desired concentration to achieve a 1:10 (0.6 × 10^4^/mL), 1:5 (1.2 × 10^4^/mL) and/or 1:2 (3.0 × 10^4^/mL) E/T ratio. All conditions were prepared in triplicate with three control wells included per experiment (target cells +/- peptide, without T cells). All outer wells were free of cells, instead containing 200 μL PBS per well.

Following addition of the primary CD8+ T cells to the target cells, the cells were placed in an incubator fitted with an Incucyte live cell imaging microscope (Sartorius). Starting 1 hour after incubator placement, four images were taken per well every hour for 48h-96h depending on the experiment. Target cell killing was subsequently assessed by comparing the confluency of the target cells in the control wells (without CD8+ T cells or with untransduced CD8+ T cells) to the confluency of the target cells in the wells with transduced primary CD8+ T cells. Analysis was conducted using the Sartorius Live Cell Analysis Software computing the target cell confluency per condition by averaging all wells within a triplicate group.

### Surface Plasmon Resonance

The DNA sequence encoding the human MHC-I was obtained from IPD-IMGT/HLA (Acc no. 142800), and the sequence encoding the β_2_M was obtained from UniProt (Acc no. P61769). The SCT construct consists of the same HA signal peptide sequence as the soluble TCRs, immediately followed by the peptide presented by the MHC-I (MAGE-A3 or Titin). The β_2_M domain is directly linked to the C-terminus of the peptide through a flexible (G_4_S)_3_ linker (L1), and to the MHC-I heavy chain via a comparatively longer (G_4_S)_4_ linker (L2). To accommodate the L1 linker, the SCT features a Y84A mutation. As with the TCR constructs, the SCTs contain a C-terminal AviTag^TM^ and 6xHis-tag for biotinylation and purification of the recombinant proteins, respectively. The recombinant SCT constructs have also been designed for cloning into the pD649 plasmid using Gibson Assembly, facilitating transfection into Expi293F cells as previously described.

SPR analysis was conducted using a BIAcore S200 system (Cytiva), using HBS-P^+^ (0.1 M HEPES, 1.5 M NaCl and 0.5% v/v Surfactant P20, pH 7.4 at 10x dilution) as a running buffer. Biotinylated MAGE-A3-A*01:01 and Titin-A*01:01 SCT complexes were immobilised onto two flow cells of Series S Sensor Chip CAP (according to the instructions in the Biotin CAPture kit, Cytiva). Between 150 and 250 RUs were immobilised in separate experiments. An irrelevant protein was also immobilised at similar levels in one flow cell to act as a control surface. Equilibrium binding analyses of TCR/p-A*01:01 interactions were carried out at 25°C by measuring binding responses at equilibrium of serial injections (twofold dilutions) of soluble non-biotinylated, 3C-treated TCR variants (a3 WT, a3a, v8 and v9), from high to low concentration, at a flow rate of 20 μl min^−1^. Biacore Evaluation Software 3.2.1 was used to conduct curve fittings and derive the equilibrium dissociation constant, *K*_D_. Experiments were repeated three times, varying the order of injected TCRs onto the chip.

### Expression and purification of the MAGE-A3 HLA-A*01:01 complex for XRC Peptide synthesis

Peptides were synthesised using Fmoc (9-fluorenylmethoxy carbonyl) solid-phase peptide synthesis and purified to >95% purity using HPLC (Genscript USA). Peptides were dissolved in dimethyl sulfoxide (DMSO) at a concentration of 200 mM and stored at -20 °C.

### HLA-A1 expression, refolding and purification

*Escherichia coli* One Shot BL21 (DE3) pLysS competent bacterial cells (Invitrogen) were used to express Human β2-microglobulin (β_2_m) and HLA-A*01:01 heavy chain residues 1-276 as inclusion bodies. Inclusion body purification was then followed using a previously described method^5,6^ involving sonication, Triton-based buffer homogenisation, and finally solubilisation in an 8M urea-MES pH 6.5 buffer at 10.0 mg/mL, which were aliquoted and stored at -80 °C.

For refolding of HLA-A*01:01 peptide complexes, a buffer containing 400 mM L-arginine monohydrochloride, 5 mM reduced glutathione, 0.5 mM oxidised glutathione, and 100 mM Tris (pH 8.0) (all from Sigma-Aldrich), supplemented with 2 mM EDTA pH 8.0 (Invitrogen) in Milli-Q water, was equilibrated at 4°C. β_2_m was added to a final concentration of 2 μM, followed after 30 minutes by addition of peptide to 15 μM. The HLA-A*01:01 heavy chain was then immediately pulsed into the refolding buffer to a final concentration of 1.5 μM.

After incubation for 72 h at 4 °C, refolded samples were passed through a 1.0 μm cellulose nitrate membrane (Cytiva) and concentrated using a VivaFlow 50R system with a 10 kDa molecular weight cut-off (Sartorius), followed by further concentration with a 10 kDa cut-off Vivaspin Turbo ultrafiltration centrifugal device (Sartorius). The concentrated material was then subjected to size exclusion chromatography on a Superdex S75 16/60 column (Cytiva) using an ÄKTA Start fast protein liquid chromatography system, eluting with 20 mM Tris pH 8.0, 150 mM NaCl. Protein-containing fractions were pooled and concentrated again using a 10 kDa cut-off Vivaspin Turbo device (Sartorius).

### Protein crystallisation

Purified v9 TCR or A3A TCR samples were mixed with MAGE-A3 in HLA-A*01:01 at equimolar ratios, at a final concentration of 10-13 mg/ml. The complexes were incubated for 30 min at 20 °C, and aggregates were subsequently removed using a centrifugal filter with a 0.22 um pore size. Sitting drop vapour diffusion crystallisation trials were set up in 96-well SwisSci plates using a Cartesian Technologies robot^32^. Drops were set up in 1:2, 1:1 and 2:1 (protein:reservoir) volume ratios, with total volumes of 200-300 nl.

Crystals of the v9 TCR/pMHC complex appeared at 15% w/v Polyethylene Glycol 3350, 0.1 M bis-Tris Propane pH 6.9, 0.2 M Potassium Nitrate, grown at 4 °C. Crystals were flash frozen in liquid nitrogen in the presence of reservoir solution supplemented with 30% (v/v) ethylene glycol as cryoprotectant.

Crystals of the A3A TCR/pMHC complex appeared at 10% w/v Polyethylene Glycol 3350, 0.1 M bis-Tris Propane pH 6.6, 0.2 M Potassium Nitrate, grown at 4 °C. Crystals were flash frozen in liquid nitrogen in the presence of reservoir solution supplemented with 30% (v/v) ethylene glycol as cryoprotectant.

### Data collection, crystallographic refinement and model analysis

X-ray diffraction data were obtained at Diamond Light Source I04, at a wavelength of 0.9537 Å. v9 TCR/pMHC crystals diffracted to 2.42 Å in space group P1. Data were indexed, integrated, and scaled using the automated xia2-DIALS^33–35^ pipeline, using the following versions XIA2 (3.23.0), DIALS (3.23.0) and CCP4 (9.0.005). All data collection and refinement statistics are summarised in Supplementary Table 1. The structure was determined by molecular replacement in Phaser (2.8.3)^36^, using the crystal structure of the MAG-IC3 TCR (PDB ID 5BRZ) as search model.

A3A TCR/pMHC crystals diffracted to 2.83 Å in space group C 1 2 1. Data were indexed, integrated, and scaled using the automated xia2 3dii pipeline^33–35^, using the following versions: XIA2 (3.21.1), DIALS (3.21) and CCP4 (9.0.005). The structure was determined by molecular replacement in Phaser (2.8.3)^36^, using the crystal structure of the MAG-IC3 TCR (PDB ID 5BRZ) as search model.

Model building was carried out by iterative rounds of manual building in COOT (0.9.8.95)^37^ and maximum-likelihood refinement in PHENIX (1.21.2)^38^, using automated X-ray and atomic displacement parameter (ADP) weight optimisation. Structure validation was performed with PHENIX using MolProbity^39^ routines.

Interface analysis was performed using the Arpeggio server^40^. TCR-pMHC engagement angles were calculated using the 3DTCR server^41^. Molecular representations were made using the program PyMOL. Calculations of root-mean-square deviations (r.m.s.d.) between structural model coordinates were done in PyMOL (The **PyMOL** Molecular Graphics System, Version 1.7 (Schrödinger, LLC, 2015)).

### Chord diagrams

Interactions for each structure were generated using Arpeggio (pdbe-arpeggio)^40^. A python script was used to compare the interactions for v9 and A3A structures and visualisation of the interaction maps and the symmetric difference between them was performed using the Matplotlib, Numpy and Pandas python libraries.

### Molecular Dynamics

#### All-Atom Molecular Dynamics Simulations

Crystal structures of A3A and v9 TCR/pMHC systems were prepared by removing non-protein residues, crystallographic waters, and ions. Structures were protonated at pH 7.4 using the H++ server^42^. Systems were prepared using tLeap and simulated with AmberTools25^43^. employing the ff19SB force field^44^ and the OPC3 water model^45^. Disulphide bonds were manually defined between appropriate cysteine residues based on crystallographic data. Each system was neutralised with counter-ions and solvated in truncated octahedral water boxes with a 10.0 Å buffer distance from the protein surface to box edges. Systems were then equilibrated to physiological ionic strength (150 mM) by adding randomised sodium and chloride ions.

The equilibration protocol consisted of several stages. Initial energy minimisation was performed for 1,000 cycles using steepest descent (50 cycles) followed by conjugate gradient optimisation, with 100 kcal mol⁻¹ Å⁻² harmonic restraints applied to all protein heavy atoms and a 10 Å non-bonded cutoff. Systems were then gradually heated from 100 K to 298 K over 1 ns using canonical ensemble (NVT) dynamics with Langevin temperature regulation (collision frequency of 1.0 ps⁻¹), while maintaining 100 kcal mol⁻¹ Å⁻² restraints on all protein heavy atoms. The initial temperature of 100 K was chosen to allow gentle heating while avoiding structural distortions during the early equilibration phase. Isothermal-isobaric ensemble (NPT) equilibration at 298 K and 1 atm followed using a Monte Carlo barostat. An initial two-stage NPT equilibration was performed with restraints applied to all protein heavy atoms, with restraint forces decreasing from 100 to 10 kcal mol⁻¹ Å⁻² across sequential 1 ns simulation. A subsequent energy minimisation (10,000 cycles; 50 steepest descent followed by conjugate gradient) with 10 kcal mol⁻¹ Å⁻² backbone restraints (Cα, C, N) was then performed. Progressive backbone-only restraint reduction was then applied through sequential 1 ns NPT simulations with restraint forces decreasing from 10 to 1 to 0.1 kcal mol⁻¹ Å⁻², followed by a final 1 ns completely unrestrained NPT equilibration. The non-bonded cutoff was set to 8 Å for all equilibration stages following initial minimisation.

Production simulations consisted of four independent 500 ns replicates for each system (A3A-pMHC, v9-pMHC, A3Aunbound, and v9 unbound) under NPT conditions at 300 K and 1atm, with pressure regulated using Monte Carlo barostat. All simulations employed a 2fs integration timestep with SHAKE constraints applied to all bonds involving hydrogen atoms. Temperature was maintained using a Langevin thermostat with a 1.0 ps^-1^collision frequency, while pressure was regulated using a Monte Carlo barostat. Long-range electrostatic interactions were treated using the Particle Mesh Ewald methods with a 9 Å non-bonded cutoff. Periodic boundary conditions were applied in all directions. Trajectory coordinated were recorded every 10ps, resulting in 50, 000 frames per 500 ns replicate, with restart filed generated every 100 ps to ensure simulation recovery.

#### RMSD & RMSF

Molecular dynamics trajectory analyses were performed using CPPTRAJ version 6.0^46^ from AmberTools25. The trajectories were centred on the MHC heavy chain with periodic boundary corrections and water/ions removed. Backbone RMSD fitting to initial frames enabled structural comparisons across simulation time for the TCR/pMHC complexes, with RMSF values quantifying residue-level flexibility.

#### Conformational Analysis

a. **Feature Extraction and Dimensionality reduction** Conformational dynamics of the CDR loops were characterised by backbone dihedral angles (φ, ψ) extracted using the CPPTRAJ multidihedral function. Residue ranges were defined according to IMGT numbering conventions using ANARCII^47^. To identify the slowest conformation degrees of freedom, we performed time-lagged independent component analysis (tICA) on the combined dataset of A3A and V9 TCRs (unbound and pMHC-bound). A lag time of 50 steps (1ns) was employed, and the top five tICS were retained.
b. **Clustering and Landscape Comparison** The resulting tICA space was discretised using k-means clustering, with the optimal number of microstates determined by maximising the VAMP-2 score. This shared clustering approach allowed for a direct comparison of the conformational landscape between systems. To quantify the divergence between A3A and V9 TCR landscapes, Bhattacharyya overlap coefficients were calculated, providing a metric of overall conformational similarity. All MSMs were validated using Chapman-Kolmogorov tests, which compared model predictions of transition probabilities at multiple lag times to direct MD measurements. Good agreement (predicted within error bars of observed) confirms that the Markov assumption holds. All analyses and visualisations were formed using the PyEMMA 2 package^48^.

#### Contact Analysis

We quantified intermolecular contacts between TCR CDR loops and pMHC using sidechain heavy atoms with a 4.5 Å distance cutoff. Contacts were tracked throughout trajectories and aggregated to the residue-pair level, calculating the percentage of simulation time each pair maintained interactions. We applied a 25% persistence threshold to identify stable, functionally relevant contacts while excluding transient fluctuations. Contact analysis examined three categories, CDR-pMHC-α1/α2 helices contacts representing primary antigen recognition and intra-TCR contacts between CDR loops and framework regions revealing internal structural binding networks.

#### MD Data Availability

All-atom MD simulation data for the A3A and A3A-pMHC TCR systems, along with associated metadata are available at https://doi.org/10.5281/zenodo.19072340. All-atom MD simulation data for the v9 and v9-pMHC TCR systems, along with associated metadata, are available at https://doi.org/10.5281/zenodo.19075350.

#### Plasmids

pYAL (Addgene)

pD649 (#156543)

pHR-SIN-CSGW (#58244)

#### Antibodies

c-Myc: Clone 9B11 AF647 Cell Signalling Tech RRID: 2233S

Streptavidin: PE BioLegend RRID: 405204

Zombie Violet™ Fixable Viability Kit: BioLegend RRID: 423114

CD8: Clone SK1 PB BioLegend RRID: 344717

αβTCR: Clone IP26 PE BioLegend RRID: 306708

CD45: Clone 2D1 PE BioLegend RRID: 368510

CD69: Clone FN50 AF647 BioLegend RRID: 310910

#### Yeast selections

autoMACSR Running Buffer - MACSR Separation Buffer: Miltenyi Biotec RRID: 130-091-221

Streptavidin Microbeads: Miltenyi Biotec RRID: 130-048-102

LS columns: Miltenyi Biotec RRID: 130-042-401

Magnetic stand: Miltenyi Biotec RRID: 130-042-303

E Buffer: 0.6 g Tris base, 91.09 g Sorbitol (1 M), 73.50 mg CaCl2 (1 mM)

Selective growth medium: SDCAA pH 4.5

Selective induction medium: SGCAA pH 4.5

Freezing medium: low-dextrose SDCAA (5 g/L dextrose)

#### Cell culture

Suspension cells were cultured in cRPMI: RPMI-1640 (Gibco) supplemented with 10% (v/v) Foetal bovine serum (Gibco), 10mM Hepes (Sigma), 1mM Sodium Pyruvate (Sigma), 1mM GlutaMax (Sigma) and antibiotics (Penicillin/Streptomycin; Sigma).

Adherent cells were cultured in: cDMEM: DMEM (Sigma), 10% (v/v) of Foetal bovine serum (Gibco; cat no. A5256801), 1mM Sodium Pyruvate (Sigma), GlutaMax (Sigma), and antibiotics (Pen/Strep; Sigma).

#### Sequences

**iTriMix19 Forward oligo** 5’ -> 3’ TTCTGTTATTGCTAGCGTTTTAGCA/iTriMix19//iTriMix19/GAK/iTriMix19//iTriMix19//iTriMi x19/iTriMix19/iTriMix19/TAYGGCGGTGGAGGTTCAGGTGGAGGTGGTAGCGGAGGCGGT GGCTCC

**pYal Reverse oligo** 5’ -> 3’ TCCACCACCACCCAAGTCTTCTTC

## Acknowledgements

**General:** We thank V. Junghans, M. Rei, J. Cabezas Caballero, T. Malinauskas, A. Gowda, O. Dushek, and T. Elliott for reagents, protocols and/or helpful discussions. We thank the staff at Diamond Light Source, T. Walter and K. Harlos for crystallisation technical support. We thank J. Rossjohn and D. R. Littler for helpful advice and discussions regarding the structural characterisation of the TCR/pMHC complexes. We thank the NHS Blood and Transplantation service at the John Radcliffe Hospital, Oxford, for donor blood samples.

## Author contributions

R.A.F. conceived of the project. J.V.M. and R.A.F. designed the overall experimental strategy and wrote the manuscript with input from A.C.H., and K.I. J.V.M. and R.A.F. edited the manuscript. J.V.M. and R.A.F. designed the A3A TCR variants. J.V.M., R.C., and A.C.H. performed protein expression and protein purification of recombinant TCRs. J.V.M. performed all yeast-display pMHC screens. J.V.M., D.H. and Y.Y. performed extraction and analysis of yeast-display deep sequencing data. M.F. designed a pipeline for quantifying peptide similarity and produced all chemical similarity heatmaps. D.H. designed pipelines for extracting deep sequencing reads and predicting wild-type agonist peptides. J.V.M. performed lentivirus production, transduction of TCRs into Jurkat T cells, and activation assays. J.V.M. performed human primary T cells isolation, transduction, and Incucyte assays. S.D.D. and A.P-F. provided Incucyte training. J.V.M. performed SPR experiments. J.V.M., R.C., and M.N.Q. performed re-folding and purification of pMHC. J.V.M. and A.C.H. set up crystallography trays. A.C.H. designed all crystallography experiments and solved the structures. C.J.T. provided structural analysis. K.I. and M.C. performed all molecular dynamics experiments. A.G.W., G.M.G., C.M.D., and R.A.F. supervised the research.

## Data and code availability

All data are available in the main text or the supplementary materials. All code will be published on GitHub prior to submission. Structure factors and coordinates for the v9 TCR- and A3A TCR pMHC complex crystal structures are deposited in the Protein Data Bank (PDB codes 29IT and 29IU, respectively).

**Supplementary Information** is available for this paper (Supplementary Tables 1 to 6).

**Correspondence and requests for materials** should be addressed to.

